# Quartet metabolite reference materials for assessing inter-laboratory reliability and data integration of metabolomic profiling

**DOI:** 10.1101/2022.11.01.514762

**Authors:** Naixin Zhang, Peipei Zhang, Qiaochu Chen, Kejun Zhou, Yaqing Liu, Haiyan Wang, Yongming Xie, Luyao Ren, Wanwan Hou, Jingcheng Yang, Ying Yu, Yuanting Zheng, Leming Shi

## Abstract

Various laboratory-developed metabolomic methods lead to big challenges in inter-laboratory comparability and effective integration of diverse datasets. As part of the Quartet Project, we established a publicly available suite of four metabolite reference materials derived from B-lymphoblastoid cell lines from a family quartet of parents and monozygotic twin daughters. We generated comprehensive LC-MS based metabolomic data from the Quartet reference materials using targeted and untargeted strategies in different laboratories. High variabilities in the qualitative and quantitative metabolomic measurements were observed across laboratories. Moreover, the Quartet multi-sample-based quality metrics were developed for objectively assessing the reliability of metabolomic profiling in detecting intrinsic biological differences among difference groups of samples. Importantly, the ratio-based metabolomic profiling, by scaling the absolute values of a study sample relative to those of a universal reference sample, enables data integration in long-term measurements across difference laboratories or platforms. Thus, we constructed the ratio-based high-confidence reference datasets between two reference samples, providing “ground truth” for inter-laboratory proficiency test, which enables objective assessment of various metabolomic methods. Our study provided the community with rich resources and best practice for objective assessment of inter-laboratory measurements and data integration, ensuring reliable large-scale and longitudinal metabolomic profiling.

## Introduction

Metabolomics is a powerful tool to discover biomarkers distinguishing biological differences in metabolite abundances related to disease diagnosis, prognosis and treatment effects^1,2^. However, the differences among such biological states are generally subtle and influenced by technical variations introduced by instruments and processing procedures^3–7^. Moreover, in large metabolomics cohort studies, batch effects are inevitable when integrating multiple batches of datasets from long-term measurement or collaboration among multiple laboratories^5,8–10^. Thus, it is crucial to assure the reliability of each batch of metabolomics measurement, as well as the integration of multiple batches of data in long-term or cross-laboratory studies so that the real signals (biological differences) can be distinguished from technical noises (unwanted variations)^11–14^.

Publicly available reference materials (RMs) are indispensable for inter-laboratory reliability assessment of current practices^15–21^. RMs in large quantities are suitable to distribute for community-wide use with the advantages of homogeneity, long-term stability, and availability of corresponding reference datasets^12^. At present, metabolite RMs have been mainly developed and distributed by the U.S. National Institute of Standards and Technology (NIST), involving many biospecimen types such as plasma, serum, urine, and liver^22–24^. By providing various types of RMs, as well as reference material suites from multiple biological states, the coverage of metabolites in the reference dataset has been improved, making it possible to compare and assess the reliability of data based on research objectives^25,26^. However, there is no renewable metabolomics reference materials suite from cultured cell lines, which represent an indispensable sample type in metabolomics studies.

Quality control (QC) metrics for objective performance evaluation are critically important. Reproducibility is one of the most widely used QC metrics, exemplified by correlations or coefficient of variation^27,28^. It helps to assess the level of unwanted variations introduced by the sample processing and detection procedures through repeated measurements of a universal reference material^29^. However, a high reproducibility from repeated measurements of a single sample does not guarantee a high resolution in identifying inherent biological differences (*i.e*., signals) among various sample groups. Identification of differentially expressed metabolites and development of predictive models to classy different sample groups are the two major goals for quantitative metabolomics technologies. Therefore, QC metrics pertinent to such research purposes are crucial to measure the ability to discriminate intrinsic biological differences among multiple sample groups^30,31^. Accuracy is another important QC metric, which is assessed through comparison of the measured metabolite concentrations with the “ground truth” in the reference datasets^25^. However, it is unachievable for evaluating the accuracy of untargeted metabolomic profiling, wherein the quantitatively measured values are usually calculated as relative output of instrumental response, which is notoriously incomparable between batches, protocols, instruments, or laboratories. Objective assessment of quantification accuracy of untargeted metabolomics is essential to ensure the reliable detection of biological differences in clinical biomarker discovery. Therefore, it is crucial to develop quality metrics to objectively evaluate the reproducibility and accuracy of metabolomics datasets at the level of detecting biological differences despite the choice of measurement strategies^32^.

Reliable integration of large-scale metabolomic data is a prerequisite for robust biomarker discovery and validation. Even if the intra-batch data is of high quality, batch effects are everywhere in large-scale metabolomics studies. In-house QC samples are widely used in long-term measurement within a single laboratory. Profiling QC samples along with study samples helps to assess the stability of measurement in each batch, and to ensure efficient integration of multiple batches by removing batch effects introduced by unwanted variations over a time span^27,33–38^. A pooled QC sample in the form of a mixture of the study samples has been widely used in this scenario, but it failed to ensure reliable data integration, mainly because the “pooled QC sample” is not identical across studies or across laboratories ^35,39,40^ and the one QC sample based metrics are not pertinent to research purposes as mentioned above. Therefore, there is a lack of best practice for objective assessment of data integration using reference materials^32^, which may hinder the cross-batch, cross-laboratory, and cross-study data integration for exploring new biological insights.

As part of the Quartet Project (chinese-quartet.org) to provide “ground truth” as well as best practices for the quality control and data integration of multiomics profiling, we established the publicly available Quartet metabolite RMs and reference datasets. The Quartet metabolite RMs enabled the research purpose related QC metric, *i.e*., the multi-sample based signal-to-noise ratio (SNR), for assessing the ability of discriminating the inherent biological differences among sample groups. In addition, we also demonstrated that the ratio-based metabolomic profiling using universal reference material(s) can enable the long-term and cross-laboratory data integration in large scale and multi-center metabolomics studies.

## Results

### Overview of the study design

In this study, we aim to provide the community with multi-sample based metabolite reference materials (RMs) suite and reference datasets for the inter-laboratory reliability assessment of metabolomic profiling using a wide range of analytical techniques. The Quartet metabolite RMs were prepared as part of the Quartet Project in which matched reference materials of DNA, RNA, proteins, and metabolites were simultaneously manufactured from the same batch of cultured cells. Four immortalized B-lymphoblastoid cell lines were derived from a Chinese Quartet family including father (F7), mother (M8), and their monozygotic twin daughters (D5 and D6) (**Fig. 1a**). The cellular metabolites were extracted using methanol: water (6:1) solution (**Extended Data Fig. 1**). Eleven external controls were added into the cellular extracts at known amounts. These spike-ins include ten flora metabolites and one xenobiotic compound, sulfadimethoxine (**Extended Data Table 1**). The cellular extracts were aliquoted into 1000 vials per cell line and then vacuum frozen dried. Each vial of the Quartet metabolite RM contains dried cellular metabolites extracted from approximately 10^6^ cells, which are suitable for most LC-MS/MS based metabolomic profiling. Additionally, the metabolite RMs were formulated as dried cellular extracts, so they cannot be used for QC of sample extraction steps.

**Fig. 1.**
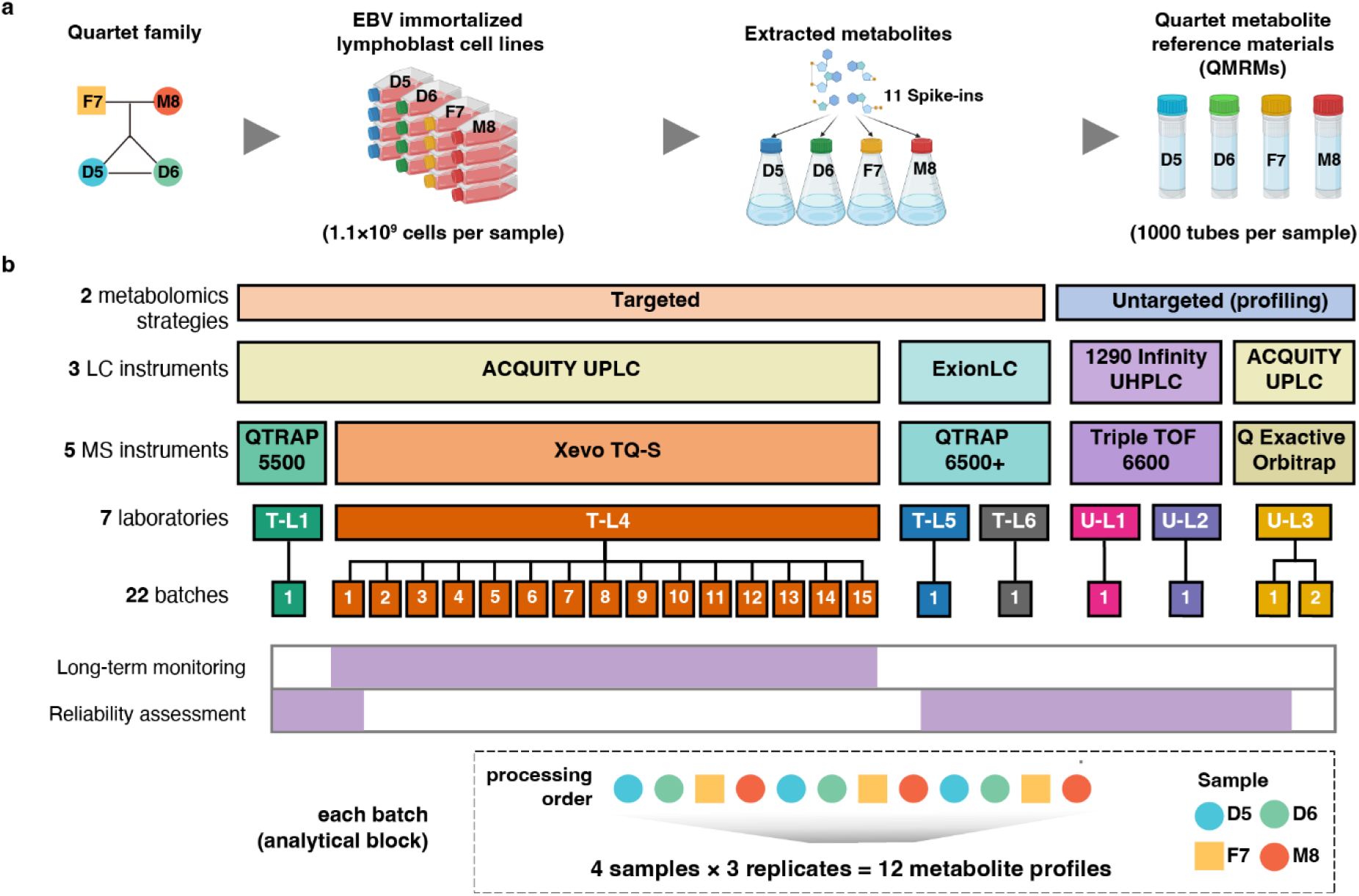
Study overview. **a**, Preparation of the Quartet metabolite reference materials. Four B-lymphoblastoid cell lines (LCLs) of a family quartet including Father (F7), Mother (M8), and monozygotic twin daughters (D5 and D6) were used for extracting metabolites. Eleven spike-ins were added to the cell extract and aliquoted into 1,000 tubes per-sample. **b**, Data generation. LC-MS based targeted (T-) and untargeted (U-) metabolomic datasets were generated in different laboratories for inter-laboratory reliability assessment. Long-term monitoring was conducted using untargeted strategy within a laboratory (T-L4) for two years.

For inter-laboratory reliability assessment of metabolomic profiling, we generated multi-laboratory datasets using untargeted and targeted strategies (**Fig. 1b**). Three replicates of each Quartet sample were measured within a batch in six laboratories. In each laboratory, the metabolomic methods have been developed independently using different liquid chromatograph (LC) and mass spectrometer (MS) instruments, which is the current practice in the field of metabolomic profiling (**Extended Data Table 2**). One laboratory 4 (T-L4) used a targeted metabolomic method to calculate the metabolite concentration with standard calibration curves, and for other targeted metabolomic methods the quantification was performed by relative metabolite abundance detected by multiple reaction monitoring (MRM). For the untargeted metabolomic methods, quantification was accomplished by relative metabolite abundance detected by precursor ions. For long-term stability monitoring of the Quartet metabolite RMs, three replicates of each Quartet sample were measured monthly in T-L4 for two years. Overall, 180 metabolomic profiles were collected.

### High variabilities in the qualitative and quantitative metabolomic measurements

We evaluated the qualitative and quantitative performance of metabolomics using the Quartet reference materials in different laboratories. High variabilities were found in the qualitative and quantitative metabolomic measurements.

The number of metabolites detected by each laboratory varied considerably, ranging from 79 (T-L1) to 462 (T-L5, **Fig. 2a**). Untargeted metabolomic strategies are usually regarded as a tool for profiling all metabolites present in a sample. However, there was no obvious advantage in the number of detected metabolites using untargeted strategies. For example, the number of detected metabolites using untargeted profiling in U-L1 was only 204, whereas the targeted strategy in T-L5 detected the largest number of metabolites, up to 462. We also compared the number of detected metabolites using different filtering criteria. Coefficient of variance (CV) was used to evaluate the reproducibility of measurement from technical replicates of the same single sample, whereas intraclass correlation coefficient (ICC) was applied to measure the reliability of metabolites from aspects of reproducibility and discriminability of multiple samples in test-retest analysis. We defined the reproducibly and reliably detectable metabolites with combined criteria of CV < 30% and ICC > 0.4, and the percentages ranged from 36% to 90%. A laboratory using untargeted strategies (U-L2) detected 402 metabolites, but only 36% of them were detected reproducibly and reliably (**Fig. 2a**). On the other side, although another laboratory using untargeted strategies (U-L3) detected 304 metabolites, 59% of these metabolites were recognized as reliable (**Fig. 2a**).

**Fig. 2.**
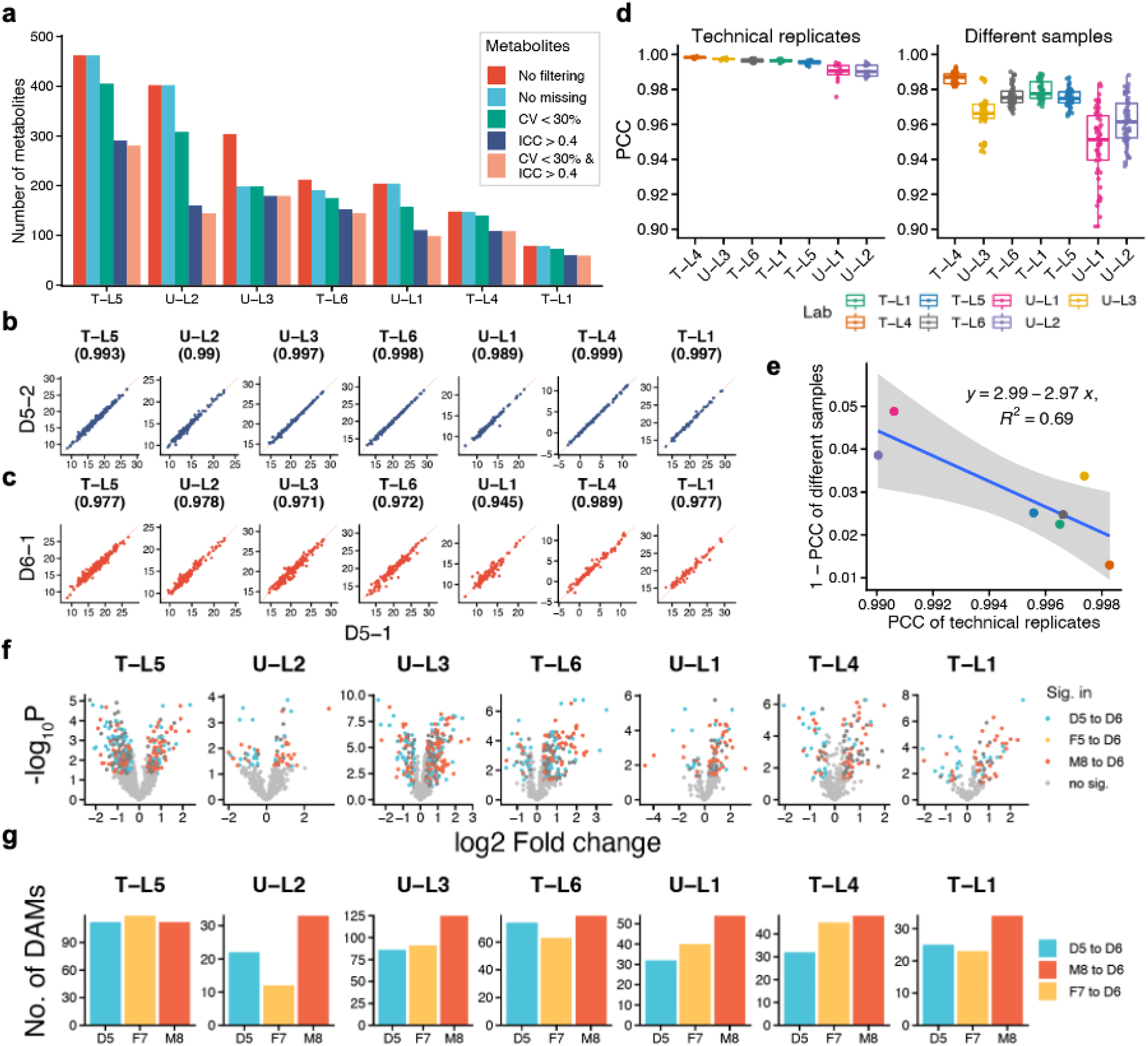
High variabilities in the qualitative and quantitative metabolomic measurements. **a**, Numbers of detected metabolites in each metabolomic measurement using different filtering criteria, including no filtering (all detected metabolites in any of the samples), no missing (metabolites detected in all 12 samples); CV (Coefficient of variance) < 30%; ICC (intraclass correlation coefficient) > 0.04; CV < 30% & ICC > 0.04. **b**, Reprehensive scatter plots of technical replicates (D5-1 and D5-2). **c**, Reprehensive scatter plots of different samples (D5-1 and D6-1). **d**, Pearson correlation coefficient (PCC) of pairs of technical replicates and of different Quartet samples in each measurement; **e**, Negative correlation between reproducibility (PCC of technical replicates) and discriminability (1-PCC of different samples). **f**, Differential abundance metabolites (DAMs) analysis for three sample pairs. Volcano plots were used to display the magnitude of the fold change versus the statistical significance level in each measurement. **g**, Numbers of DAMs identified for three sample pairs in each measurement.

Pearson correlation coefficient (PCC) of pairs of technical replicates indicated the reproducibility of quantitative profiles, while PCC of pairs of different Quartet metabolite RMs indicated the level of discriminability of biological differences. As shown in **Fig. 2b**, the PCC of technical replicates of D5 was high in each of the seven datasets, ranging from 0.989 to 0.999. However, the PPC between D5 and D6 was also high, ranging from 0.945 to 0.989 (**Fig. 2c**). For example, the PCC of technical replicates in T-L4 was 0.999, indicating high reproducibility. However, the PCC between different samples in the same laboratory was 0.989, indicating low discriminability. These results indicated that a high reproducibility of technical replicates does not guarantee a high resolution in identifying inherent biological differences (discriminability) between different sample groups (**Fig. 2d**). In addition, there was a negative correlation between PCC of technical replicates and (1-PCC) of difference sample groups (**Fig. 2e**).

Identification of differential abundance metabolites (DAMs) is a major goal of biomarker discovery using metabolomic technologies. However, as shown in **Fig. 2f** where the volcano plots were used to display the magnitude of the fold change versus the statistical significance level, big differences in both the fold changes and statistical significance levels were seen when comparing the same sample pairs. The number of DAMs ranged from ~10 to ~120 in D5/D6, F7/D6, and M8/D6 comparisons (**Fig. 2g**). In two of the laboratories (T_L5 and U_L3) that identified the highest number of DAMs, most of the DAMs were more highly expressed in D6 than in D5, F7, and M8 at T-L5. However, most of the DAMs were more highly expressed in D5, F7, and M8 than in D6 at U-L3. Taken together, these results implicated that achieving high inter-laboratory reproducibility of DAMs was challenging.

### Inter-laboratory reliability assessment by Quartet based signal-to-noise ratio

Based on the Quartet multi-sample design, we designed a signal-to-noise ratio (SNR) metric to take reproducibility and discriminability into consideration simultaneously. SNR is calculated as the ratio of the averaged distance between different Quartet samples (“signal”) to the averaged distances between technical replicates for each sample (“noise”) on a 2D-PCA scatter plot (**Fig. 3c**)^41,43^, where a higher SNR indicates better reproducibility and discriminability (**Fig. 3**). As expected, for a measurement to be considered reliable, the sample-to-sample difference should be bigger than variation of technical replicates.

**Fig. 3.**
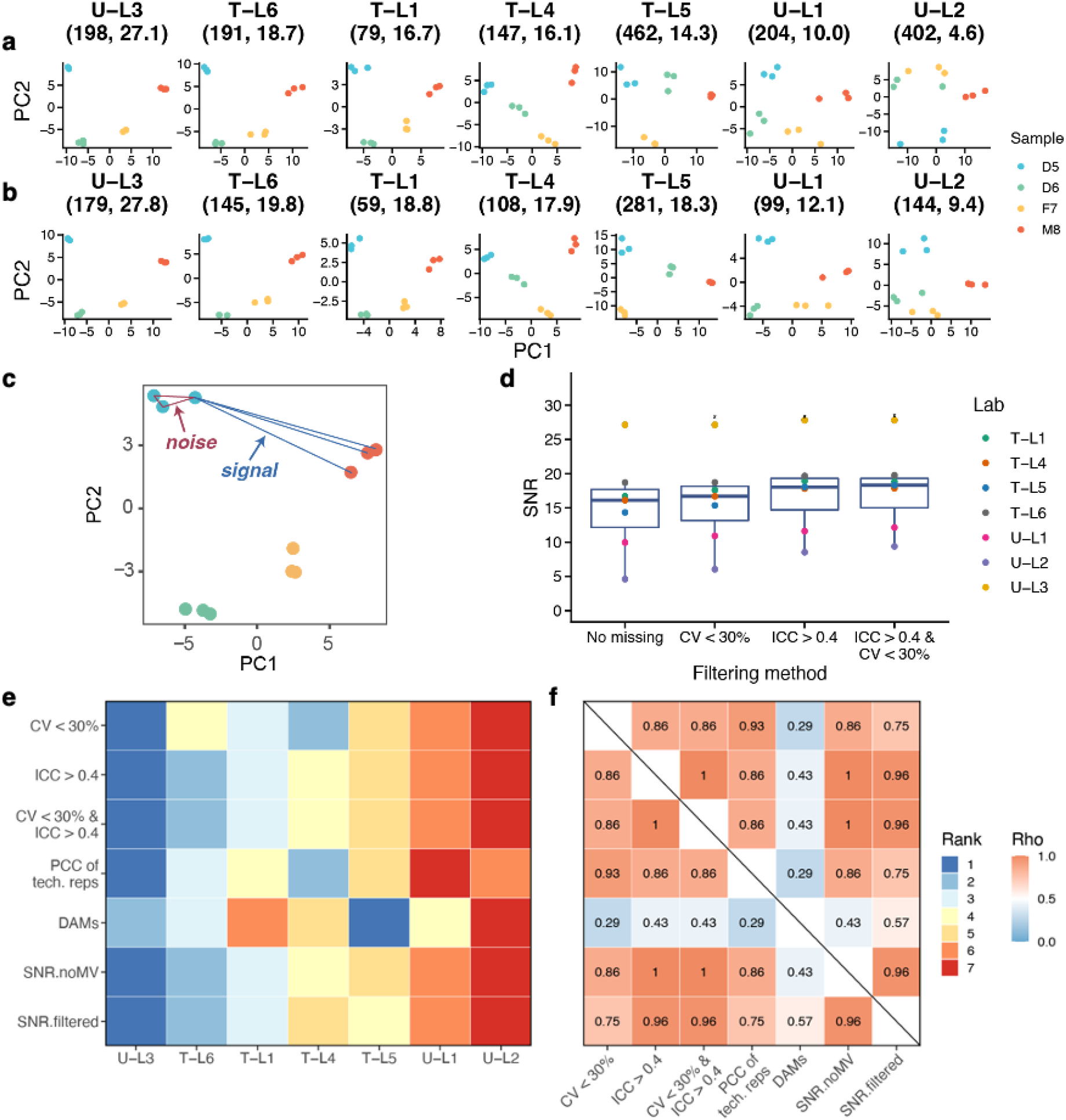
Inter-laboratory reliability assessment by Quartet based Signal-to-Noise ratio. **a** and **b**, Reliability assessment using signal-to-noise ratio (SNR) in different laboratories with (**a**) or without (**b**) filtration. The results were visualized by PCA plots. The number of features used and the calculated SNR were shown above each plot. **c**, Schematic diagram of SNR calculated as the ratio of the averaged distance between different Quartet samples (“signal”) to the averaged distances between technical replicates for each sample (“noise”) on a 2D-PCA scatter plot. **d**, SNRs calculated with metabolites filtered with different criteria. **e**, The inter-laboratory data quality was ranked using different QC metrics, including the percentages of retained metabolites using different filtering criteria (CV<30%; ICC>0.4; CV<30% and ICC>0.4), PCC of technical replicates, the number of DAMs, and SNR calculated with or without filtering. **f**, The concordances of data quality ranking using different QC metrics.

We computed the SNR for each batch of metabolomic profiling (4 sample × 3 replicates) using metabolites detected in all 12 samples in the batch. As shown in the PCA, the first two principal components demonstrated clear separation among the four reference samples in good-quality metabolomic profiling data, but not in poor-quality data (**Fig. 3a**). Astonishingly, high variabilities in data quality were observed in these metabolomic datasets (range 4.6~27.1). After filtering with combined criteria (CV<30% and ICC>0.4) to retain the reliably detectable metabolites, the SNRs were slightly improved but the relative quality ranking of batches did not change obviously (**Fig. 3b**). **Fig. 3d** illustrated the SNRs calculated with metabolites filtered with different criteria, indicating the robustness of SNRs in evaluating the laboratory-specific reliability with or without filtering metabolites.

We also ranked the quality of the metabolomic datasets generated in different laboratories using different QC metrics, including the percentages of retained metabolites using different filtering criteria (CV<30%; ICC>0.4; CV<30% and ICC>0.4), PCC of technical replicates, the numbers of DAMs, and SNR calculated with or without filtering. As shown in **Fig. 3e**, the inter-laboratory data quality rankings were not entirely concordant using different QC metrics. T-L5 outperformed others in terms of the numbers of discovered DAMs, but did not perform well by other QC metrics. These results implicated that more DAMs did not guarantee good data quality, which can be confound by false positive discovery. In addition, T-L4 performed well by PCC of technical replicates, but did not perform well by SNR or DAMs. These results suggested that correlation of replicates from one reference material did not have enough resolution in identifying the among-sample differences.

The overall concordances among these QC results were also evaluated in **Fig. 3f**. QC results assessed by DAMs showed low concordance with all the other results, and the percentages of retained features after filtering (CV<30% and ICC>0.4) was highly correlated with SNR. These data suggested that the inter-laboratory data quality assessments were dependent on the QC metrics being used. PCC of technical replicates from one reference material and the number of DAMs between two reference materials did not show enough resolution in metabolomic profiling quality in identifying differences among sample groups. The Quartet multi-sample based SNR provided an objective QC metric for inter-laboratory reliability assessment for a wide range of metabolomic technologies.

### Ratio-based metabolite profiling enables data integration across laboratories

To evaluate the reliability of metabolomic data integration, we examined the qualitative and quantitative performance of the integrated data generated in different laboratories. We first evaluated the qualitative concordance of detected metabolites among these datasets. However, most detected metabolites were reported by only one laboratory, and merely six metabolites were reported in all the seven metabolomic datasets (**Fig. 4a**). The intersection size of detected metabolites among different laboratories were shown in **Extended Data Fig. 2**, and there were only 58 metabolites detected by all the three global metabolomic profiling strategies. These results demonstrated the poor concordance of metabolite identification across different laboratories.

**Fig. 4.**
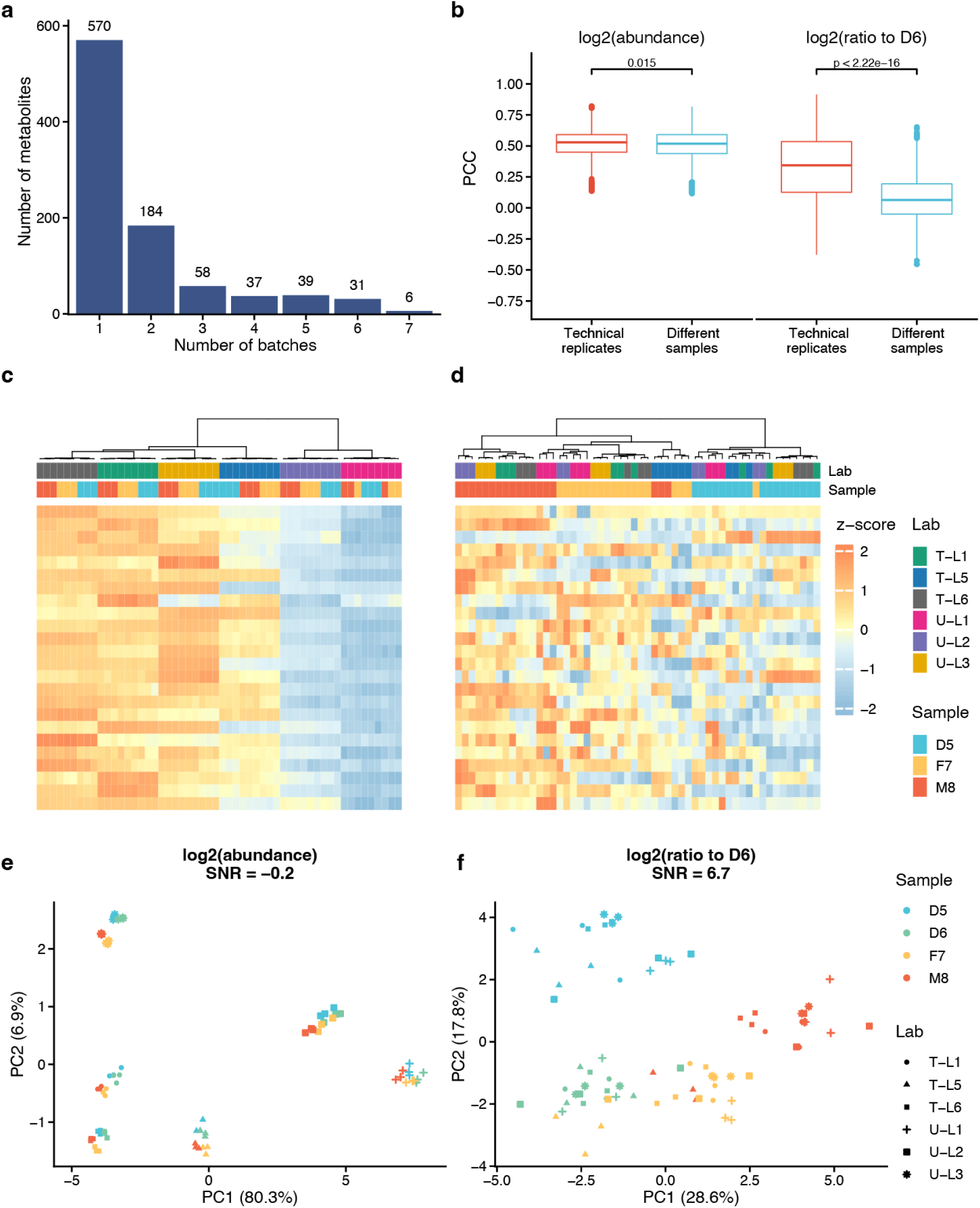
Ratio-based metabolite profiling enables data integration across laboratories. **a**, Qualitative concordance of metabolite identification. The numbers of metabolites detected in different batches of metabolomic datasets were shown. **b**, Pearson correlation coefficients (PCCs) of pairs of technical replicates and of different Quartet samples were compared using quantitative datasets at abundance level or ratio to D6 level. **c** and **d**, Cross-lab data integration was visualized by hierarchical cluster analysis (HCA) at absolute abundance level (**c**) and relative ratio to D6 level (**d**). **e** and **f**, Cross-lab data integration assessment using signal-to-noise ratio (SNR) by principal component analysis (PCA) at absolute abundance level (**e**) and relative ratio to D6 level **(f**).

To further evaluate the quantitative reliability of cross-laboratory integration of metabolomic data at absolute abundance level, we first compared the differences between PCCs of technical replicates and those between different reference sample groups. However, the differences between the two types of PCCs were not dramatic (**Fig. 4b**, **Left**). Hierarchical cluster analysis (HCA) and principal component analysis (PCA) were also used to visualize the magnitude of technical variation of data integration at the absolute abundance level. The integrated cross-laboratory metabolomic profiling data were first clustered by batch (laboratory) but not by different sample groups (**Fig. 4c**). Similar results were demonstrated in PCA (**Fig. 4e**), where the first principal component (PC1) clearly showed the dramatic differences between the six batches of data and the distinct Quartet samples cannot be separated.

Importantly, after converting the absolute abundance data to a ratio scale relative to the same reference material (D6) on a metabolite-by-metabolite basis in each batch, significant differences between PCCs of technical replicates and those of different samples were found (**Fig. 4b, Right**). Similar results were supported by HCA and PCA plots (**Figs. 4d** and **4f**). After ratio-based scaling, the metabolomic profiling relative to D6 first clustered by the four different Quartet sample groups (**Fig. 4d**). PCA plots showed clear separation of the four groups of reference samples (D5, D6, F7, and M8) and the drastic batch effects seen at the absolute abundance (**Figs. 4c** and **4e**) largely disappeared.

Our results showed that batch effects were prevalent in cross-laboratory metabolomic data integration at the absolute abundance level, presenting a real challenge for large-scale integrative analyses of multi-center data. Fortunately, the cross-laboratory data reliability can be greatly improved by converting the absolute abundance to a ratio-based metabolomic profiling using universal reference materials such as the Quartet metabolite reference materials.

### Ratio-based metabolite profiling improves data integration in long-term measurement

In order to evaluate the long-term stability of metabolomic profiling, we generated a total of 15 batches of Quartet metabolomic data in T-L4 over a period of two years. This targeted metabolomic strategy calculates the concentration of each metabolite with a specific standard calibration curve, and is referred to as the “absolute” quantification approach of metabolomic profiling and regarded as one of the most reliable ones.

We first evaluated the qualitative concordance of detected metabolites in long-term measurement. We found that only 100 out of 148 metabolites (67.6%) were detected in all the 15 batches of datasets (**Fig. 5a**). Thus, the stability of metabolite identification was still not ideal even for the absolute quantification metabolomics strategy.

**Fig. 5.**
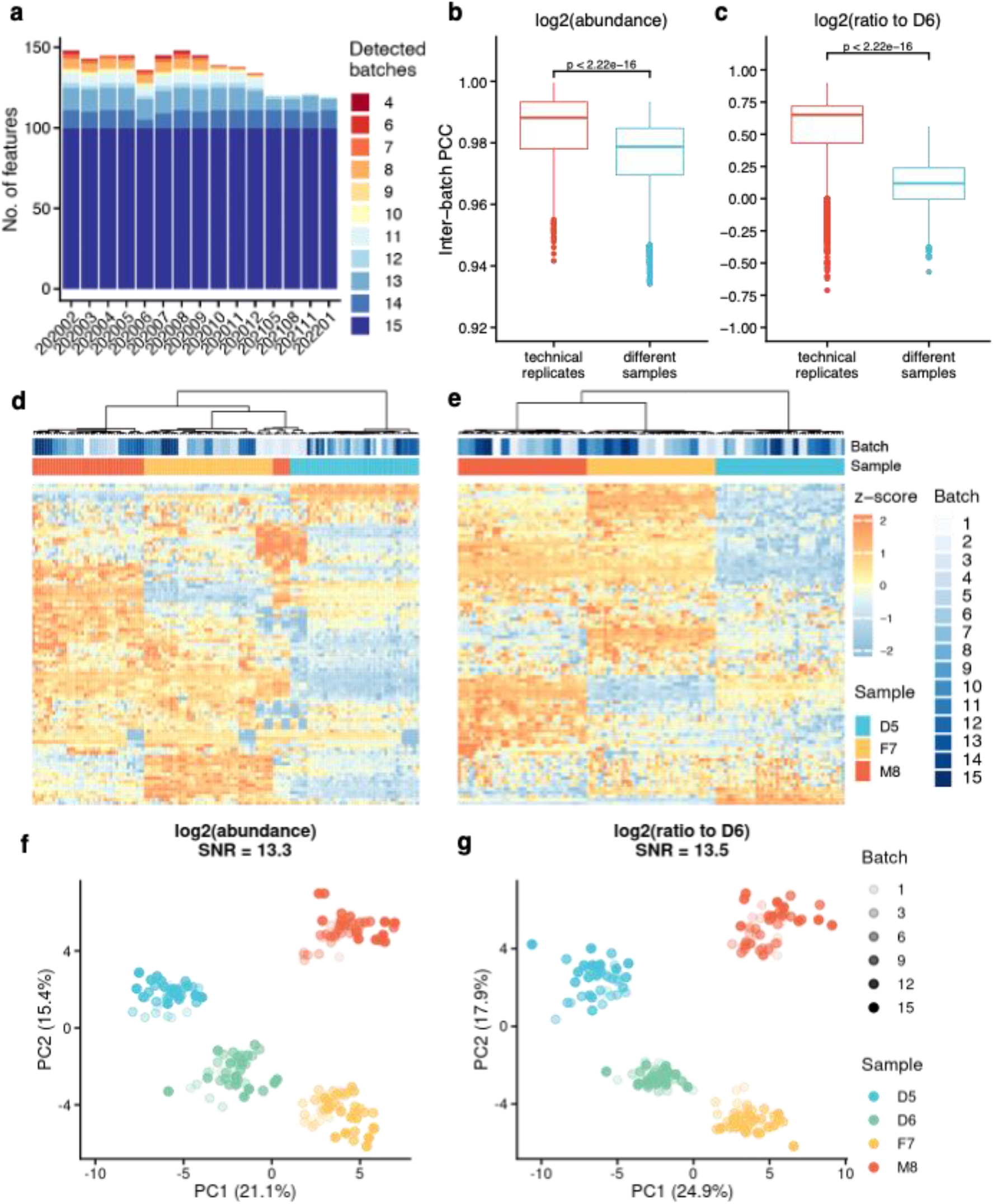
Ratio-based metabolite profiling improves data integration in long-term monitoring. **a**, Qualitative concordance of metabolite identification. The numbers of metabolites detected in each batch of metabolomic datasets were shown. **b** and **c**, Pearson correlation coefficients (PCCs) of pairs of technical replicates and of different Quartet samples were compared using quantitative datasets at abundance level (b) or ratio to D6 level (c). **d** and **e**, Cross-batch data integration was visualized by hierarchical cluster analysis (HCA) at absolute abundance level (**d**) and relative ratio to D6 level (**e**). **e** and **f**, Cross-batch data integration assessment using signal-to-noise ratio (SNR) by principal component analysis (PCA) at absolute abundance level (**f**) and relative ratio to D6 level (**g**).

To further evaluate the quantitative stability of long-term metabolomic data at the absolute concentration level, we first compared the differences between PCCs of technical replicates and those between different reference sample groups. There existed significant differences between the two types of PCCs (**Fig. 5b**). HCA and PCA were also used to visualize the magnitude of technical variation of data integration at the absolute concentration level. Most of the samples in the integrated long-term metabolomic dataset were clustered by different sample groups, but several M8 samples mis-clustered into the F7 sample group (**Figs. 5d** and **5f**). In addition, the Quartet signal-to-noise ratio was calculated to be 13.3 for the integrated dataset, indicating a good separation among different Quartet samples.

We also integrated the long-term metabolomic datasets by converting the absolute concentration values to a ratio scale relative to those of the same reference material (D6) on a metabolite-by-metabolite basis per batch. The difference between PCCs of technical replicates and PCCs of different samples, a surrogate of discriminability, increased dramatically from 0.009 (absolute, **Fig. 5b**) to 0.532 (relative, **Fig. 5c**). Similar results were observed by HCA and PCA plots (**Figs. 5e** and **5g**). After ratio-based scaling, all the samples clustered correctly by different Quartet sample groups (**Fig. 5e**). The Quartet signal-to-noise ratio was improved slightly from 13.3 to 13.5 (**Fig. 5g**). Using the Levey-Jennings plot, we continuously monitored each metabolite measurement across runs (**Extended Data Fig. 3**). There were 57, 14, and 10 metabolites deviated from the mean beyond ±3 SD for 3 RMs (D5, F7, M8), demonstrating evidence of systematic errors (**Extended Data Fig. 3, up**). After ratio-based scaling to D6 sample, the number of systematically deviated metabolites decreased to 8, 11, and 15 for D5, F7, and M8, respectively (**Extended Data Fig. 3, down**).

These results showed that the reliability of long-term metabolomic profiling measurement can still be improved by converting the absolute concentration values to a ratio scale using universal reference materials such as the Quartet metabolite reference materials.

### Construction of ratio-based Quartet metabolite reference datasets

In order to provide “ground truth” reference datasets for evaluating the accuracy of metabolomic quantification, we constructed the ratio-based metabolomic reference datasets by scaling the absolute abundance values of D5, F7, and M8 relative to those of D6. **Fig. 6a** illustrated the workflow of consensus integration from seven metabolomics datasets from different laboratories. First, the metabolites detected in all three replicates in each dataset were defined as detected metabolites, and the union of the detected metabolites in all the seven datasets were 939, 944, 948, and 948 for the four reference materials (D5, D6, F7, and M8). Secondly, 210 reproducibly detectable metabolites were retained in all the four reference samples in more than one dataset. Thirdly, ratio-based values were calculated using differential abundance metabolites (DAMs) for each sample pair (D5/D6, F7/D6, or M8/D6) with *p* < 0.05 in more than one dataset. Finally, the high-confidence ratio-based reference value for each metabolite is defined as a geometric mean of fold-changes calculated from each replicate of more than one dataset. After these filtrations, the first release of the Quartet ratio-based metabolomic datasets (v.1.0) for each sample pair (D5/D6, F7/D6, and M8/D6) contained 47, 44, and 51 high-confidence metabolites, respectively (**Extended Data Tables 3–5**).

**Fig. 6.**
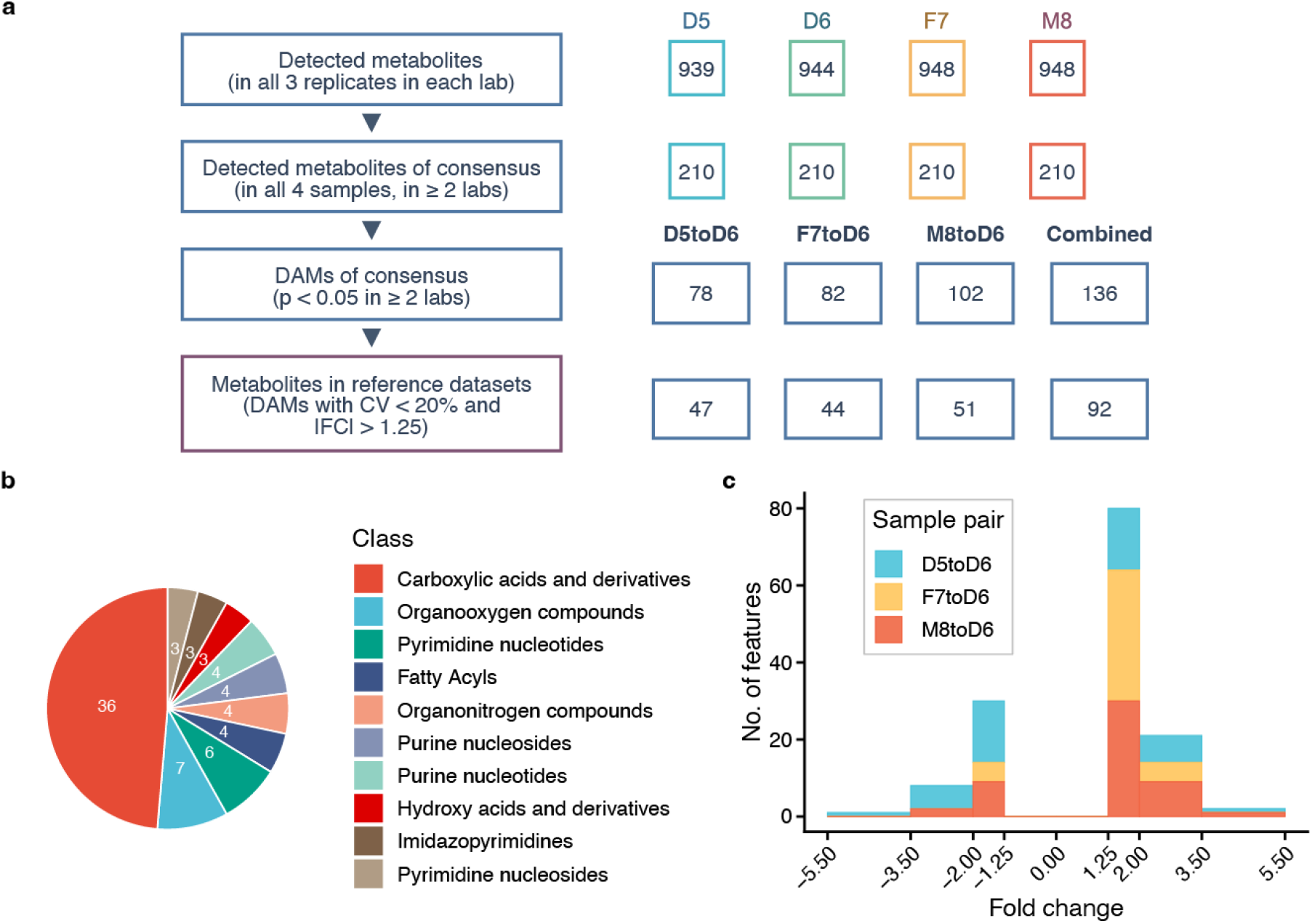
Integration of ratio-based Quartet metabolite reference datasets. **a**, The workflow of integration of ratio-based metabolite reference datasets. **b**, The number of metabolites in high-confidence reference datasets annotated into 10 classes according to the HMDB database. **b**, The distribution histogram of fold changes of the high-confidence reference metabolites for three sample pairs.

The union of high-confidence reference metabolites for the three sample pairs were annotated into 10 classes according to the HMDB database (https://hmdb.ca) (**Fig. 6b**). The most abundant class was carboxylic acids and derivatives, with 36 metabolites contained in the reference datasets. The ratio values for the high-confidence reference metabolites for each sample pair were summarized in **Fig. 6c**, covering a wide range of fold-changes from −4.4 to 5.1. With the advance in metabolomic technologies and the generation of additional datasets, the Quartet metabolomic reference datasets will be updated periodically through the Quartet Data Portal^46^ (http://chinese-quartet.org).

### Best practice for inter-laboratory proficiency test of metabolomic profiling using the Quartet metabolite reference materials

Inter-laboratory proficiency test is essential to achieve reliable metabolomic profiling. We recommend profiling the Quartet reference materials (*e.g*., four samples × three replicates) for method validation. We provided two types of QC metrics for quality assessment of quantitative metabolomic datasets. One is the Quartet multi-sample based SNR to measure the ability to discriminate the intrinsic biological differences among different reference samples. The other is the quantitative concordance between the evaluated batch of data and the reference datasets, calculated by relative correlation (RC). The recall of detected DAMs was recommended to qualitatively assess the accuracy against the Quartet reference datasets (**Fig. 7a**).

**Fig. 7.**
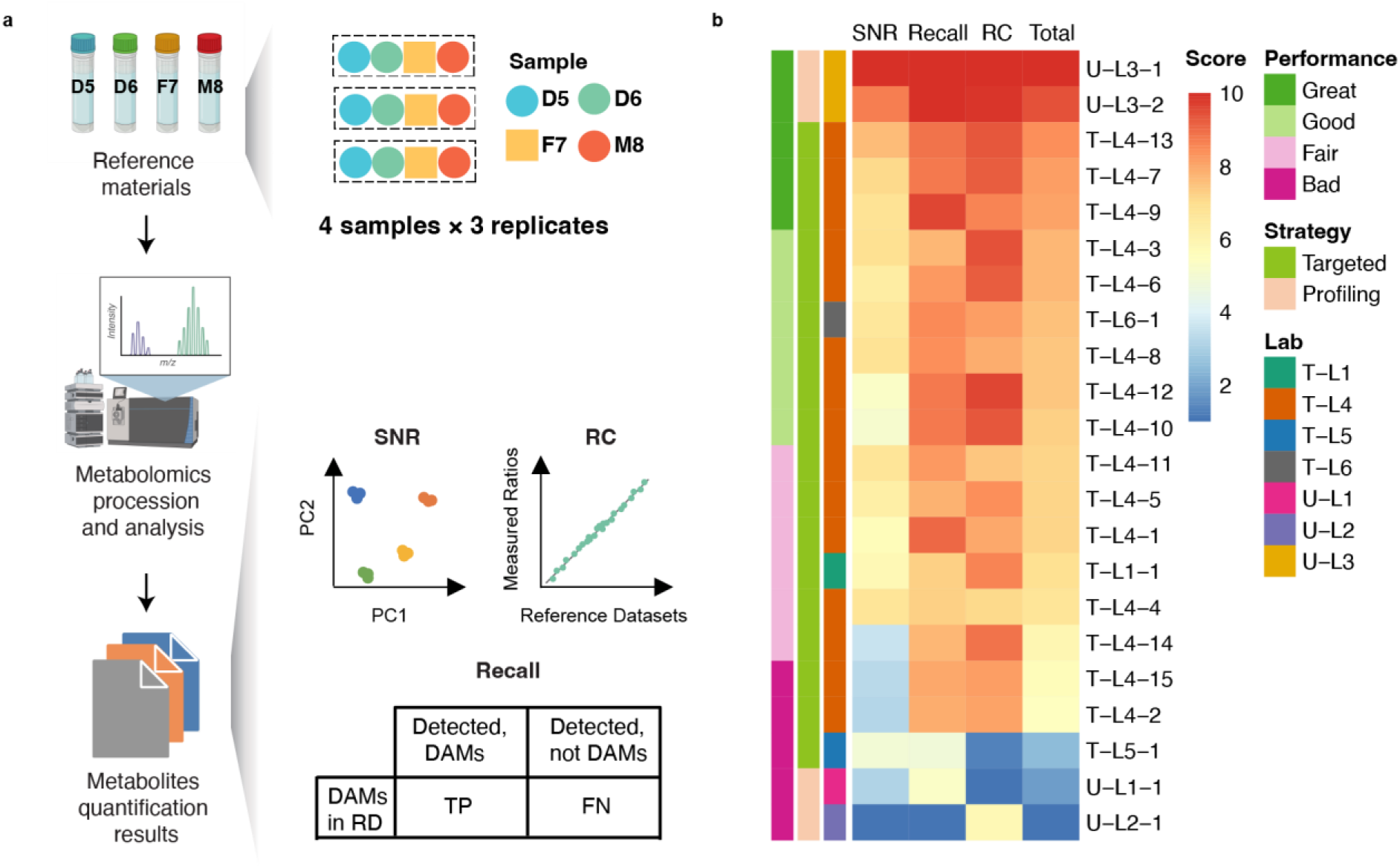
Best practice for inter-laboratory proficiency test of metabolomic profiling using Quartet metabolite reference materials. **a**, Flowchart of an inter-laboratory proficiency test using the Quartet metabolite reference materials. **b**, Inter-laboratory proficiency for each of the 22 batches of metabolomic datasets using targeted or untargeted strategies with SNR, RC, and recall. The overall performance was classified into four levels (Great, Good, Fair, and Bad)

We evaluated the laboratory proficiency for each of the 22 batches of metabolomic datasets using targeted or untargeted strategies with SNR, RC, and recall. As shown in **Fig. 7b**, the relative quality rankings using different QC metrics were not concordant. The correlations among the three metric values were relatively low (**Extended Data Fig. 4**), therefore a total score would be better to rank the quality of metabolomic datasets. Using the total score calculated using SNR, RC, and recall with each metric value scaled from 0 to 10, we found that the inter-laboratory proficiency was not dependent on the metabolomic strategy. The top one laboratory proficiency was achieved in U_L3 using an untargeted strategy. The three datasets with the worst laboratory proficiency (U-L1, T-L5, and U-L2) were generated in three laboratories using either untargeted or targeted strategies. These results supported the notion that the Quartet metabolite reference materials and related QC metrics were suitable for a wide range of metabolomic technologies using both targeted and untargeted strategies. In addition, the standardized QC workflow for inter-laboratory comparisons can be performed through the Quartet Data Portal^46^, where the relative quality ranking among the cumulative metabolomic datasets can be obtained.

## Discussion

As part of the Quartet project^41^, we provide the community the first suite of renewable metabolite reference materials, with matched DNA^42^, RNA^43^, and protein^44^ isolated from the same immortalized cell lines. The intended use of the Quartet metabolite reference materials includes intra-laboratory quality control, inter-laboratory proficiency test, and quality assurance of large-scale metabolomic data generation and integration.

We also defined the ratio-based reference datasets as abundance ratios of sample pairs (D5/D6, F7/D6, and M8/D6) between the Quartet reference materials, providing “ground truth” for assessing the quantification accuracy of a wide range of metabolomic technologies. There are two clear advantages of the ratio-based reference datasets. First, the ratio-based quantification can mitigate technical variations seen at the absolute-level quantification, because the instrumental output can systematically change by different laboratory-developed methods^41,45^. Therefore, the Quartet ratio-based reference datasets are suitable for proficiency test and method validation of a wide range of targeted and non-targeted metabolomic strategies using various LC-MS instruments. Secondly, the accuracy and reproducibility of the ratio-based metabolite qualification indicate the ability of a metabolomic measurement procedure in distinguishing the biological differences among sample groups, which is a basis for biomarker discovery and validation using metabolomic technologies. Although the v1.0 of the metabolite reference datasets covers only 92 metabolites, the reference datasets will be updated periodically through the Quartet Data Portal (http://chinese-quartet.org/) community-wide participation^46^. Our study suggested a paradigm shift of defining the ground truth for assessing the accuracy of metabolomic profiling.

The Quartet multi-sample based SNR is developed as an objective QC metric for assessing the ability in identifying intrinsic biological differences between various groups of samples, which is the basis for the development and application of metabolomic profiling based clinical diagnostics. Compared with the previously widely used one-sample based technical reproducibility for quality assessment of quantitative profiling, the unique Quartet SNR guaranteed a high resolution in identifying serious quality issues. An obvious limitation of any “ground truth” dependent QC metrics is due to their reliance on easily detectable metabolites. However, the Quartet based SNR is applicable for assessing the reliability of global metabolomic profiling without considering the ground truth, making it a complementary and broadly applicable approach for objective quality assessment.

Using the Quartet metabolite reference materials for inter-laboratory proficiency testing, high variabilities of the qualitative and quantitative metabolomic measurements were observed across laboratories. We found that the best and worst performing batches were all generated using the untargeted metabolomics strategies. This result suggested that the intrinsic laboratory proficiency was critically important for developing in-house untargeted metabolomics strategies, consistent with a previous report^47^. In addition, even for an “absolute” quantification method using a targeted strategy that quantifies each metabolite by standard calibration curves within the same laboratory, clear technical variations were observed by continuously measuring the same Quartet reference materials over a long period of time. This result implicated that profiling reference materials in each batch along with the study samples help monitor long-term data quality and technical drift within a laboratory.

Importantly, the multi-sample based SNR can also be used to objectively monitor and correct batch effects. To achieve reliable data integration from long-term and cross-laboratory large-scale metabolomics profiling, we recommend using universal reference materials per-batch along with study samples. As long as the integrated datasets maintain the ability to differentiate the different Quartet samples, the reliability of the metabolomic data from the study samples for further exploratory metabolomic biomarker discovery is assured.

Our study also demonstrated the potential utility of the Quartet reference materials as a universal reference sample to scale the absolute abundance to ratio-based metabolomic profiling. The ratio-based metabolomic profiling was suitable for internal quality control in longitudinal measurement within a laboratory, and it can also be used to calibrate the metabolomic profiling in multiple centers. Even if the metabolomic methods were developed using different wet-lab operation procedures on different LC-MS instruments, the ratio-based metabolomic data integration was reliable enough for differentiating the various Quartet reference materials. The intrinsic batch-effect resistant characteristics of the ratio-based profiling is also demonstrated by other quantitative omics profiling technologies, such as methylomics, transcriptomics, and proteomics^41, 43, 45^.

Although we demonstrated the importance of using the Quartet metabolite reference materials and corresponding QC metrics in ensuring reliable biological discovery, there are some limitations beyond the scope of this study. First, the Quartet metabolite reference materials were extracted cellular metabolites in the form of lyophilized power and could not be applied to in the QC of the sample preparation procedures. Moreover, the metabolites extracted from cells could not fully cover metabolites from other sources of biospecimen, such as plasma, serum and tumor tissues, which may hinder the wider application of the Quartet reference materials especially when the matrix of the study samples is largely different from cellular extractants. However, with the Quartet multi-sample design, the reference data dependent and independent QC metrics could be used to comprehensively assess the system-specific reliability of biological discoveries.

In summary, as an important part of the Quartet multiomics reference materials suites consisting DNA, RNA, proteins, and metabolites, the Quartet metabolite reference materials, the reference datasets, and the corresponding quality metrics help lay the foundation for reliable discovery of metabolomic differences through quality control of the intra- and inter-laboratory data generation and integration processes.

## Methods

### Ethics approval and consent to participate

The study was approved by the IRB (Institutional Review Board) of the School of Life Sciences, Fudan University (BE2050) and conformed to the principles set out in the 1975 Declaration of Helsinki. Written informed consent to participate in multiomics research and allow collection of biospecimens was approved by the IRB and obtained from the Quartet family that includes monozygotic twin daughters (D5 and D6) and their parents (father, F7 and mother, M8) in Taizhou, Jiangsu Province, China.

### Quartet immortalized B-lymphoblastoid cell lines

Quartet immortalized B-lymphoblastoid cell lines were established through the infection with Epstein-Barr virus (EBV)^48^ and culturing using the protocols described in the Quartet main paper^41^. Briefly, immortalized lymphoblastoid cell lines were obtained by isolating peripheral blood mononuclear cells (PBMCs), sorting naive B cells and infecting with EBV by centrifugation at 2000 rpm for 1 hour. Lymphoblastoid cell lines were cultured in RPMI 1640 supplemented with 15% of non-inactivated FBS and 1% Penicillin-Streptomycin. Flasks were incubated on the horizontally position at 37°C under 5% CO_2_. Cell cultures were split every three days for maintenance as described in the literature^49^. Cells growing in suspension were centrifuged at 300 g for 5 min to obtain cell pellets and were washed twice with cold PBS, then store at −80°C. About 1×10^11^ cells were harvested for each cell line in the same batch to ensure that matched multiomics reference materials were extracted from the same batch of cultured cells. About 1.1×10^9^ cells per cell line were used for generating Quartet metabolite reference materials. In all cases, all cell lines were handled in parallel using exactly the same reagents and equipment, and experiments were initiated at the same time-point of the day.

### Metabolite extraction

Metabolites were extracted from EBV immortalized lymphoblastoid cell lines in L4 (Laboratory 4). At first, we thawed cells (11 tubes per sample, 1×10^8^ cells per tube) slowly on ice-bath to minimize potential sample degradation, and then added 2.4 mL ice cold methanol solution (methanol: water = 6:1) to each tube of samples. Then, the ice water bath was under ultrasonic treatment for 3 times, each time for 3 s, with an interval of 2 minutes, power 10%. Then we found that the cell mass at the bottom of the tube was completely broken by ultrasound and appeared as white emulsion. Finally, after centrifugation at 4500 g, 4°C, 20 minutes (Allegra X-15R, Beckman Coulter, Inc., Indianapolis, IN, USA), we transferred supernatant containing extracted metabolites to a new centrifuge tube.

### Quartet metabolite reference materials

Eleven external controls were spiked into the supernatant at known concentrations as internal standards, including ten metabolites commonly found in plasma (Indoleacetic acid, Taurocholic acid, Glycocholic acid, Cholic acid, Tauroursodeoxycholic acid, Taurodeoxycholic acid, Glycoursodeoxycholic acid, Glycodeoxycholic acid, Ursodeoxycholic acid, and Deoxycholic acid) and one drug sulfadimethoxine (Extended Data Table 1). For each Quartet sample, supernatant containing metabolites extracted from 1.1×10^9^ cells and 11 spike-ins were aliquoted into 1,000 vials using an automated liquid handler (Biomek 4000, Beckman Coulter, Inc., Brea, California, USA), with 5 μL solution per tube. After centrifugation at 4°C and under vacuum (Labconco, Kansas City, Missouri, USA) for 50 minutes, water was removed, and we obtained the Quartet metabolite reference materials in the form of lyophilized power. Reference materials from different Quartet samples were clearly marked with differently colored dispensing caps and labels. The tube cap colors of the reference materials D5, D6, F7, and M8 are blue, green, yellow, and red, respectively. We stored the Quartet metabolite reference materials at −80°C and shipped with dry ice.

### Sample preparation

We distributed 12 vials (triplicates for each Quartet sample) of the Quartet metabolite reference materials as a batch to each laboratory and offered basic guidance on sample preparation. The same sample running order (D5-1, D6-1, F7-1, M8-1, D5-2, D6-2, F7-2, M8-2, D5-3, D6-3, F7-3, and M8-3) in each batch was maintained among all laboratories.

#### T-L1/U-L1

Samples were first centrifuged before the researchers added 200 μL acetonitrile-water (1:1, v/v) to reconstitute. The solution was then centrifuged at 14,000 rcf for 15 min at 4°C to extract supernatant for MS analysis.

#### U-L2

The researchers added 100 μL of 50% acetonitrile to reconstitute (containing isotope-labeled internal standard mixture). The solution was vortexed for 30 s, and sonicated in ice-water bath for 10 min. After centrifugation at 13000 rpm for 15 min at 4 °C, the supernatant of 70 μL was transferred into the sample bottle and tested on the machine.

#### U-L3

The researchers added 500 μL of ice-cold 80% methanol solution to dissolve the sample. Then the solution was divided into five fractions: two for analysis by two separate reverse phases (RP)/UPLC-MS/MS methods with positive ion mode electrospray ionization (ESI), one for analysis by RP/UPLC-MS/MS with negative ion mode ESI, one for analysis by HILIC/UPLC-MS/MS with negative ion mode ESI, and one sample was reserved for backup. Samples were placed briefly on a TurboVap® (Zymark) to remove the organic solvent. The sample extracts were stored overnight under nitrogen before preparation for analysis.

#### T-L4

The researchers added 350 μL of ice-cold 50% methanol solution to dilute the sample. The plate was then stored at 20°C for 20 minutes and then centrifuged at 4000g for 30 minutes at 4°C. They transferred 135 μL of supernatant to a new 96-well plate, which contained 15 μL of internal standard per well. Serial dilutions of derivatized standards were added to the left wells. The plate was sealed for LC-MS analysis.

#### T-L5

The researchers added the 500 μL of 10% methanol solution to dissolve the powder, and then injected samples into LC-MS for analysis.

#### T-L6

The researchers added 100 μL reconstituted solution (acetonitrile: water=1:1) of HPLC-grade to the 1.5 mL EP tube containing the dried metabolites, and vortexed for 1 min; centrifuged at 15,000 rpm for 10 min at 4°C (Note: the centrifuge needs to be pre-cooled); used a 200 μ L pipette to draw about 60 μL of the supernatant and transfer to the injection vial, making sure that there are no air bubbles at the bottom of the liner or the injection vial; mixed the remaining liquid in the same sample EP tube (took an equal volume) into the same 1.5 mL EP tube, centrifuged at 15,000 rpm for 10 min at 4°C, transferred the supernatant to the injection vial as QC sample (Note: the whole process needs to be operated on ice).

### Laboratory instrument

Each laboratory used different HPLC/UPLC or MS/MS platforms to detect and quantify metabolites (details in Extended Data Table 2). L1 used Waters UPLC-MRM with AB SCIEX QTRAP 5500 mass spectrometers by targeted strategy (T-L1) and Agilent UHPLC with AB SCIEX Triple TOF 6600 mass spectrometers by untargeted strategy (U-L1). L2 used HPLC with SCIEX mass spectrometers by untargeted strategy (U-L2). L3 used Waters UPLC with Thermo Fisher Q Exactive and Orbitrap mass spectrometers by untargeted strategy (U-L3). L4 used Waters UPLC with a Waters Xevo TQ-S mass spectrometer by targeted strategy (T-L4). L5 and L6 used AB SCIEX Exion UPLC-MRM with AB SCIEX QTRAP® 6500+ mass spectrometers by targeted strategy (T-L5 and T-L6).

### Data processing

Raw data acquired using UPLC-MS were pre-processed by each participating laboratory to provide structured data in .xls format for subsequent statistical analysis. Chromatography-MS data for a single sample are a matrix of m/z versus retention time (or index) versus ion current or intensity.

#### T-L1

MRM raw data were extracted by MRMAnalyzer (R), and the peak area of each metabolite was obtained. More detailed description can be found in reference^50^.

#### U-L1

The raw data was converted into mzXML format by ProteoWizard. The researchers used the XCMS program for peak alignment, retention time correction and peak area extraction. For structure identification of metabolites, accurate mass matching (<25 ppm) and secondary spectrum matching were used to search the laboratory’s inhouse-built database.

#### U-L2

The researchers used ProteoWizard software to convert the original mass spectrum into mzXML format and XCMS for retention time correction, peak identification, peak extraction, peak integration, and peak alignment. An inhouse-built secondary mass spectrometry database was used in parallel to identify the peaks.

#### U-L3

The researchers used ThermoFisher Scientific software Xcalibur QuanBrowser for peak detection and integration. A detailed description of data processing including chromatographic alignment, QC practices and compound identification can be found in reference^51^.

#### T-L4

The raw data files generated by UPLC-MS/MS were processed using the QuanMET software (v2.0, Metabo-Profile, Shanghai, China) to perform peak integration, calibration, and quantitation for each metabolite.

#### T-L5

The detection of the experimental samples using MRM (Multiple Reaction Monitoring) were based on T-L5 inhouse database. The Q3 was used for metabolite quantification. The Q1, Q3, RT (retention time), DP (declustering potential) and CE (collision energy) were used for metabolite identification. The data files generated by HPLC-MS/MS were processed using the SCIEX OS Version 1.4 to integrate and correct the peak. The main parameters were set as follows: minimum peak height, 500; signal/noise ratio, 5; Gaussian smooth width, 1. The area of each peak represents the relative content of the corresponding metabolite.

#### T-L6

The MRM raw data were extracted by OS-MQ software (AB SCIEX), and the peak area value of each metabolite was obtained.

### Data integration

We collected 264 metabolomics profiles at the metabolite level from all laboratories, with each laboratory provided HMDB (Human Metabolome Database, https://hmdb.ca) IDs corresponding to the metabolites.

We integrated these metabolomics profiles first by their HMDB IDs and then by metabolite names. Metabolites were annotated into different classes with the information downloaded from HMDB (https://hmdb.ca/system/downloads/current/hmdb_metabolites.zip, released on 2021-11-17).

### Performance metrics

Based on Quartet metabolite RMs and RDs, we constructed three types of performance metrics to comprehensively evaluate the reproducibility and accuracy of each laboratory in detecting biological differences. Among them, signal-to-noise ratio (SNR) was designed to evaluate the ability of each laboratory in extracting different Quartet samples from technical replicates. Recall of differential abundance metabolites (DAMs) and relative correlation (RC) were computed based on the reference datasets and were designed to evaluate the ability and accuracy in detection of biological differences among Quartet sample pairs.

#### Signal-to-noise ratio (SNR)

We measured SNR through comparing the average Euclidean distances between different Quartet samples (“signals”) to those between different technical replicates of the same Quartet sample(“noises”) computed based on the first two principal components of PCA, which was same to other companion articles from the Quartet multiomics project. SNR was defined as the following equation:

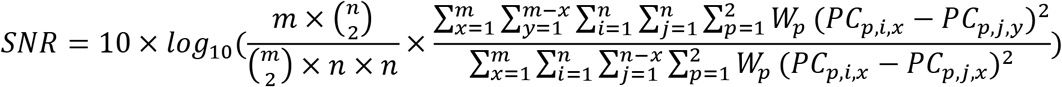

Here, *m* was the number of different groups of samples, and *n* was the number of technical replicates of the same sample group. The variances explained by the p^th^ principal component (*PC*_*p*_) was noted as *W*_*p*_. *PC*_*p*,*i*,*x*_, *PC*_*p*,*j*,*x*_ and *PC*_*p*,*j*,*y*_represent the value of i^th^ and j^th^ replicate of sample *x* or sample *y* on p^th^ principal component, respectively.

#### Recall

We computed Recall for the assessment of qualitative agreement with the RDs, as the fraction of the differential abundance metabolites (DAMs) in RDs that are successfully retrieved. Here recall is the number of measured DAMs (p < 0.05, t test) divided by the number of DAMs should be identified as RDs.

#### Relative correlation (RC)

We measured RC for the assessment of quantitative consistency with the RDs. First, we calculated the average log2 abundance of each metabolite of each Quartet sample. Based on the average log2 abundance, we computed relative abundance values of metabolites of each sample pair (log2 ratios to D6) overlapped with the RDs in each laboratory. Finally, the Pearson correlation was computed between the measured relative abundance values and consensual ones in the RDs.

### Statistical analysis

We used R version 4.0.5 and associated packages to perform all statistical analysis. All statistical tests described in this work were two-sided. Tests involving comparisons of distributions were done using ‘wilcox.test’ unless otherwise specified. Intraclass correlation coefficient (ICC) was computed based on package *irr* (v0.84.1), using two-way model and estimated by the agreement between raters to compute differences in judges’ mean ratings. We plotted all results based on R package *ggplot2* (v3.3.6), *cowplot* (v1.1.1), *ComplexUpset* (v1.3.3), *ggpubr* (v0.4.0), *ggsci* (v2.9) and *GGally* (v2.1.2).

## Materials Availability

The Quartet metabolite reference materials can be requested for research use from the Quartet Data Portal (http://chinese-quartet.org/) under the Administrative Regulations of the People’s Republic of China on Human Genetic Resources.

## Data and Code Availability

The Quartet metabolite Reference datasets could also be downloaded from the Quartet Data Portal. Metabolomics profiles generated from all laboratories included in this article could be downloaded from National Omics Data Encyclopedia (NODE project OEP000970, https://www.biosino.org/node/project/detail/OEP000970) under the regulation of the Human Genetic Resources Administration of China (HGRAC).

## Acknowledgments

This study was supported in part by National Key R&D Project of China (2018YFE0201603 and 2018YFE0201600), the National Natural Science Foundation of China (31720103909 and 32170657), Shanghai Municipal Science and Technology Major Project (2017SHZDZX01), State Key Laboratory of Genetic Engineering (SKLGE-2117), and the 111 Project (B13016). Some of the illustrations in this paper were created with BioRender.com.

## Author contributions

Y.Z., L.S., and Y.Y. conceived and oversaw the study. Y.Z., P.Z., K.Z., H.W., W.H. cultured the cell lines, prepared or characterized the metabolite reference materials. Y.Z., K.Z., and Y.X. coordinated and/or performed metabolomic data generation. N.Z., P.Z., Q.C., Y.L., L.R., J.Y., Y.Y., Y.Z., and L.S. performed data analysis and/or interpretation. J.Y. and Y.L. managed the datasets. N.Z., Y.Z., and L.S. wrote the manuscript. All authors reviewed and approved the manuscript.

## Extended data

**Extended Data Fig. 1.**
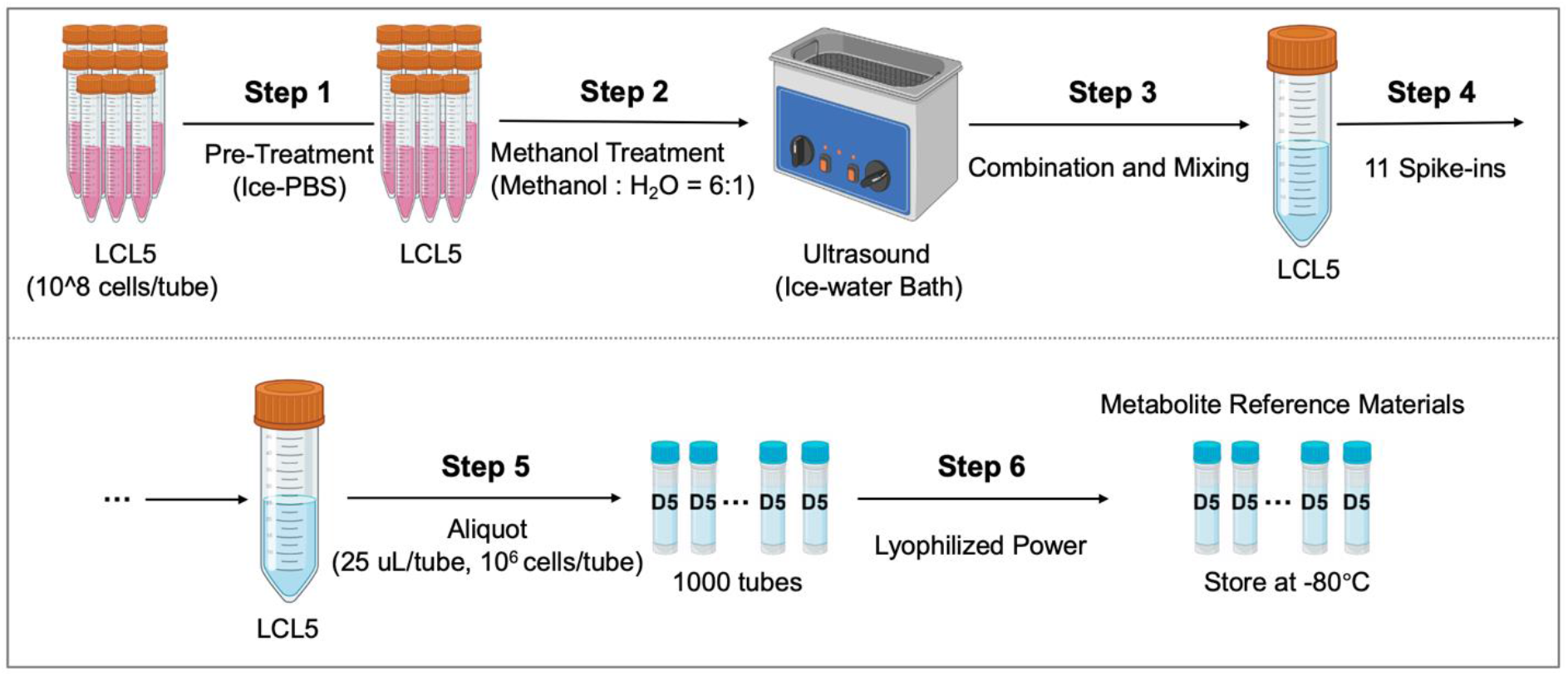
Steps of the preparation of Quartet metabolite reference materials.

**Extended Data Table 1.** External controls spiked in the Quartet metabolite reference materials.

**Extended Data Fig. 1 | Steps of the preparation of Quartet metabolite reference materials.**
| No. | External controls | Amount (pmol) | Comment |
| --- | --- | --- | --- |
| 1 | Indoleacetic acid | 25 | Intestinal bacteria metabolite |
| 2 | Taurocholic acid | 1 | Primary bile acids |
| 3 | Glycocholic acid | 5 |  |
| 4 | Cholic acid | 25 |  |
| 5 | Tauroursodeoxycholic acid | 2.5 | Secondary bile acids |
| 6 | Taurodeoxycholic acid | 7.5 |  |
| 7 | Glycoursodeoxycholic acid | 1 |  |
| 8 | Glycodeoxycholic acid | 0.5 |  |
| 9 | Ursodeoxycholic acid | 25 |  |
| 10 | Deoxycholic acid | 50 |  |
| 11 | Sulfadimethoxine | 5 | Sulfa antibiotics |

**Extended Data Table 2.** Experimental methods of metabolomics profiling in each dataset.

| Lab | Strategies | Liquid Chromatograph (LC) Instrument | Chromatographic Column | Mass Spectrometer (MS) Instrument | Metabolites Coverage |
| --- | --- | --- | --- | --- | --- |
| T-L1 | Targeted (MRM based) | Waters ACQUITY UPLC | Waters ACQUITY UPLC BEH Amide (1.7µm, 2.1 x 100 mm column) | AB SCIEX QTRAP 5500 | 79 |
| U-L1 | Untargeted | Agilent Technologies 1290 Infinity UHPLC | Waters ACQUITY UPLC BEH Amide (1.7µm, 2.1 × 100 mm column) | AB SCIEX Triple TOF 6600 | 203 |
| U-L2 | Untargeted | Agilent Technologies 1290 Infinity UHPLC | Waters ACQUITY UPLC BEH Amide (1.7µm, 2.1 x 100 mm column) | AB SCIEX Triple TOF 6600 | 400 |
| U-L3 | Untargeted | Waters ACQUITY UPLC | Waters ACQUITY UPLC BEH Amide (1.7µm, 2.1 x 150 mm column) | Thermo Fisher Q Exactive and Orbitrap | 262 |
| T-L4 | Targeted | Waters ACQUITY UPLC | Waters ACQUITY UPLC BEH C18 (1.7µM, 2.1 × 100 mm column) | Waters Xevo TQ-S | 148 |
| T-L5 | Targeted (MRM based) | AB SCIEX ExionLC | Waters ACQUITY UPLC HSS T3 (1.8µM, 2.1 × 100 mm column) and BEH C8 (1.7µm, 2.1 × 100 mm column) | AB SCIEX QTRAP 6500+ | 462 |
| T-L6 | Targeted (MRM based) | AB SCIEX ExionLC | Waters XBridge BEH Amide (3.5µm, 4.6 × 100mm column) | AB SCIEX QTRAP 6500+ | 207 |

**Extended Data Table 3.** Reference datasets of D5toD6.

| HMDB ID | Metabolite name | Fold change | KEGG ID | Class |
| --- | --- | --- | --- | --- |
| HMDB0000902 | NAD | 3.23 | C00003 | (5'→5')-dinucleotides |
| HMDB0001173 | 5'-Methylthioadenosine | 1.82 | C00170 | 5'-deoxyribonucleosides |
| HMDB0001185 | S-Adenosylmethionine | 1.57 | C00019 | 5'-deoxyribonucleosides |
| HMDB0000714 | Hippuric acid | -1.81 | C01586 | Benzene and substituted derivatives |
| HMDB0000001 | 1-Methylhistidine | 1.49 | C01152 | Carboxylic acids and derivatives |
| HMDB0000052 | Argininosuccinic acid | 2.9 | C03406 | Carboxylic acids and derivatives |
| HMDB0000092 | Dimethylglycine | -1.29 | C01026 | Carboxylic acids and derivatives |
| HMDB0000159 | L-Phenylalanine | -1.67 | C00079 | Carboxylic acids and derivatives |
| HMDB0000161 | L-Alanine | -1.37 | C00041 | Carboxylic acids and derivatives |
| HMDB0000187 | L-Serine | 1.52 | C00065 | Carboxylic acids and derivatives |
| HMDB0000191 | L-Aspartic acid | 2.21 | C00049 | Carboxylic acids and derivatives |
| HMDB0000202 | Methylmalonic acid | 1.44 | C02170 | Carboxylic acids and derivatives |
| HMDB0000214 | Ornithine | 1.97 | C00077 | Carboxylic acids and derivatives |
| HMDB0000254 | Succinic acid | 1.38 | C00042 | Carboxylic acids and derivatives |
| HMDB0000267 | Pyroglutamic acid | -1.76 | C01879 | Carboxylic acids and derivatives |
| HMDB0000517 | L-Arginine | 2.02 | C00062 | Carboxylic acids and derivatives |
| HMDB0000883 | L-Valine | -1.29 | C00183 | Carboxylic acids and derivatives |
| HMDB0000904 | Citrulline | -2.55 | C00327 | Carboxylic acids and derivatives |
| HMDB0001325 | N6,N6,N6-Trimethyl-L-lysine | 1.83 | C03793 | Carboxylic acids and derivatives |
| HMDB0001511 | Phosphocreatine | 1.48 | C02305 | Carboxylic acids and derivatives |
| HMDB0001539 | Asymmetric dimethylarginine | 2.49 | C03626 | Carboxylic acids and derivatives |
| HMDB0000300 | Uracil | -1.39 | C00106 | Diazines |
| HMDB0000086 | Glycerophosphocholine | 5.18 | C00670 | Glycerophospholipids |
| HMDB0000126 | Glycerol 3-phosphate | 2.1 | C00093 | Glycerophospholipids |
| HMDB0000190 | L-Lactic acid | -1.93 | C00186 | Hydroxy acids and derivatives |
| HMDB0001311 | D-Lactic acid | -2.62 | C00256 | Hydroxy acids and derivatives |
| HMDB0000157 | Hypoxanthine | -1.36 | C00262 | Imidazopyrimidines |
| HMDB0000292 | Xanthine | -2.4 | C00385 | Imidazopyrimidines |
| HMDB0001096 | Carbamoyl phosphate | 1.4 | C00169 | Organic phosphoric acids and derivatives |
| HMDB0000097 | Choline | -1.3 | C00114 | Organonitrogen compounds |
| HMDB0000210 | Pantothenic acid | 1.51 | C00864 | Organooxygen compounds |
| HMDB0000230 | N-Acetylneuraminic acid | -3.76 | C19910 | Organooxygen compounds |
| HMDB0000247 | Sorbitol | -2.47 | C00794 | Organooxygen compounds |
| HMDB0000613 | Erythronic acid | -1.92 |  | Organooxygen compounds |
| HMDB0000765 | Mannitol | -2.41 | C00392 | Organooxygen compounds |
| HMDB0000855 | Nicotinamide riboside | 2.09 | C03150 | Organooxygen compounds |
| HMDB0000779 | Phenyllactic acid | -1.72 | C01479 | Phenylpropanoic acids |
| HMDB0000133 | Guanosine | -1.68 | C00387 | Purine nucleosides |
| HMDB0000195 | Inosine | -1.57 | C00294 | Purine nucleosides |
| HMDB0000299 | Xanthosine | 1.62 | C01762 | Purine nucleosides |
| HMDB0000045 | Adenosine monophosphate | 1.46 | C00020 | Purine nucleotides |
| HMDB0000089 | Cytidine | -1.33 | C00475 | Pyrimidine nucleosides |
| HMDB0000296 | Uridine | -1.41 | C00299 | Pyrimidine nucleosides |
| HMDB0000788 | Orotidine | -2.54 | C01103 | Pyrimidine nucleosides |
| HMDB0000095 | Cytidine monophosphate | 1.97 | C00055 | Pyrimidine nucleotides |
| HMDB0000288 | Uridine 5'-monophosphate | 1.95 | C00105 | Pyrimidine nucleotides |
| HMDB0001564 | CDP-ethanolamine | 1.82 | C00570 | Pyrimidine nucleotides |

**Extended Data Table 4.** Reference datasets of F7toD6.

| HMDB ID | Metabolite name | Fold change | KEGG ID | Class |
| --- | --- | --- | --- | --- |
| HMDB0000902 | NAD | 1.58 | C00003 | (5'->5')-dinucleotides |
| HMDB0000462 | Allantoin | 1.42 | C01551 | Azoles |
| HMDB0000056 | beta-Alanine | 1.38 | C00099 | Carboxylic acids and derivatives |
| HMDB0000064 | Creatine | 1.33 | C00300 | Carboxylic acids and derivatives |
| HMDB0000072 | cis-Aconitic acid | 1.5 | C00417 | Carboxylic acids and derivatives |
| HMDB0000094 | Citric acid | 1.66 | C00158 | Carboxylic acids and derivatives |
| HMDB0000112 | gamma-Aminobutyric acid | -1.63 | C00334 | Carboxylic acids and derivatives |
| HMDB0000148 | L-Glutamic acid | 1.29 | C00025 | Carboxylic acids and derivatives |
| HMDB0000176 | Maleic acid | 1.37 | C01384 | Carboxylic acids and derivatives |
| HMDB0000177 | L-Histidine | -1.43 | C00135 | Carboxylic acids and derivatives |
| HMDB0000182 | L-Lysine | 1.48 | C00047 | Carboxylic acids and derivatives |
| HMDB0000202 | Methylmalonic acid | 1.77 | C02170 | Carboxylic acids and derivatives |
| HMDB0000254 | Succinic acid | 1.85 | C00042 | Carboxylic acids and derivatives |
| HMDB0000446 | N-alpha-Acetyl-L-lysine | 1.38 | C12989 | Carboxylic acids and derivatives |
| HMDB0000517 | L-Arginine | 1.37 | C00062 | Carboxylic acids and derivatives |
| HMDB0000562 | Creatinine | 1.29 | C00791 | Carboxylic acids and derivatives |
| HMDB0000766 | N-Acetyl-L-alanine | 1.31 |  | Carboxylic acids and derivatives |
| HMDB0001511 | Phosphocreatine | 1.77 | C02305 | Carboxylic acids and derivatives |
| HMDB0000235 | Thiamine | 1.78 | C00378 | Diazines |
| HMDB0000300 | Uracil | 1.36 | C00106 | Diazines |
| HMDB0000201 | L-Acetylcarnitine | -1.37 |  | Fatty Acyls |
| HMDB0000222 | Palmitoylcarnitine | -1.6 | C02990 | Fatty Acyls |
| HMDB0000086 | Glycerophosphocholine | 1.28 | C00670 | Glycerophospholipids |
| HMDB0000156 | Malic acid | 1.76 | C00149 | Hydroxy acids and derivatives |
| HMDB0000190 | L-Lactic acid | 1.58 | C00186 | Hydroxy acids and derivatives |
| HMDB0000157 | Hypoxanthine | 1.28 | C00262 | Imidazopyrimidines |
| HMDB0000292 | Xanthine | 1.58 | C00385 | Imidazopyrimidines |
| HMDB0000935 | Uridine diphosphate glucuronic acid | 2.45 | C00167 | Lactones |
| HMDB0000062 | L-Carnitine | 1.39 | C00318 | Organonitrogen compounds |
| HMDB0001257 | Spermidine | 1.63 | C00315 | Organonitrogen compounds |
| HMDB0000210 | Pantothenic acid | 1.55 | C00864 | Organooxygen compounds |
| HMDB0000613 | Erythronic acid | 1.53 |  | Organooxygen compounds |
| HMDB0000855 | Nicotinamide riboside | 3.09 | C03150 | Organooxygen compounds |
| HMDB0000217 | NADP | 1.59 | C00006 | Phenols |
| HMDB0000299 | Xanthosine | 2.22 | C01762 | Purine nucleosides |
| HMDB0000175 | Inosinic acid | 1.5 | C00130 | Purine nucleotides |
| HMDB0001554 | Xanthylic acid | 2.01 | C00655 | Purine nucleotides |
| HMDB0000229 | Nicotinamide ribotide | 1.84 | C00455 | Pyridine nucleotides |
| HMDB0000296 | Uridine | 1.38 | C00299 | Pyrimidine nucleosides |
| HMDB0000286 | Uridine diphosphate glucose | 1.49 | C00029 | Pyrimidine nucleotides |
| HMDB0000288 | Uridine 5'-monophosphate | 1.67 | C00105 | Pyrimidine nucleotides |
| HMDB0000290 | Uridine diphosphate-N-acetylglucosamine | 2.18 | C00043 | Pyrimidine nucleotides |
| HMDB0000295 | Uridine 5'-diphosphate | 1.91 | C00015 | Pyrimidine nucleotides |
| HMDB0000653 | Cholesterol sulfate | -1.32 | C18043 | Steroids and steroid derivatives |

**Extended Data Table 5.**
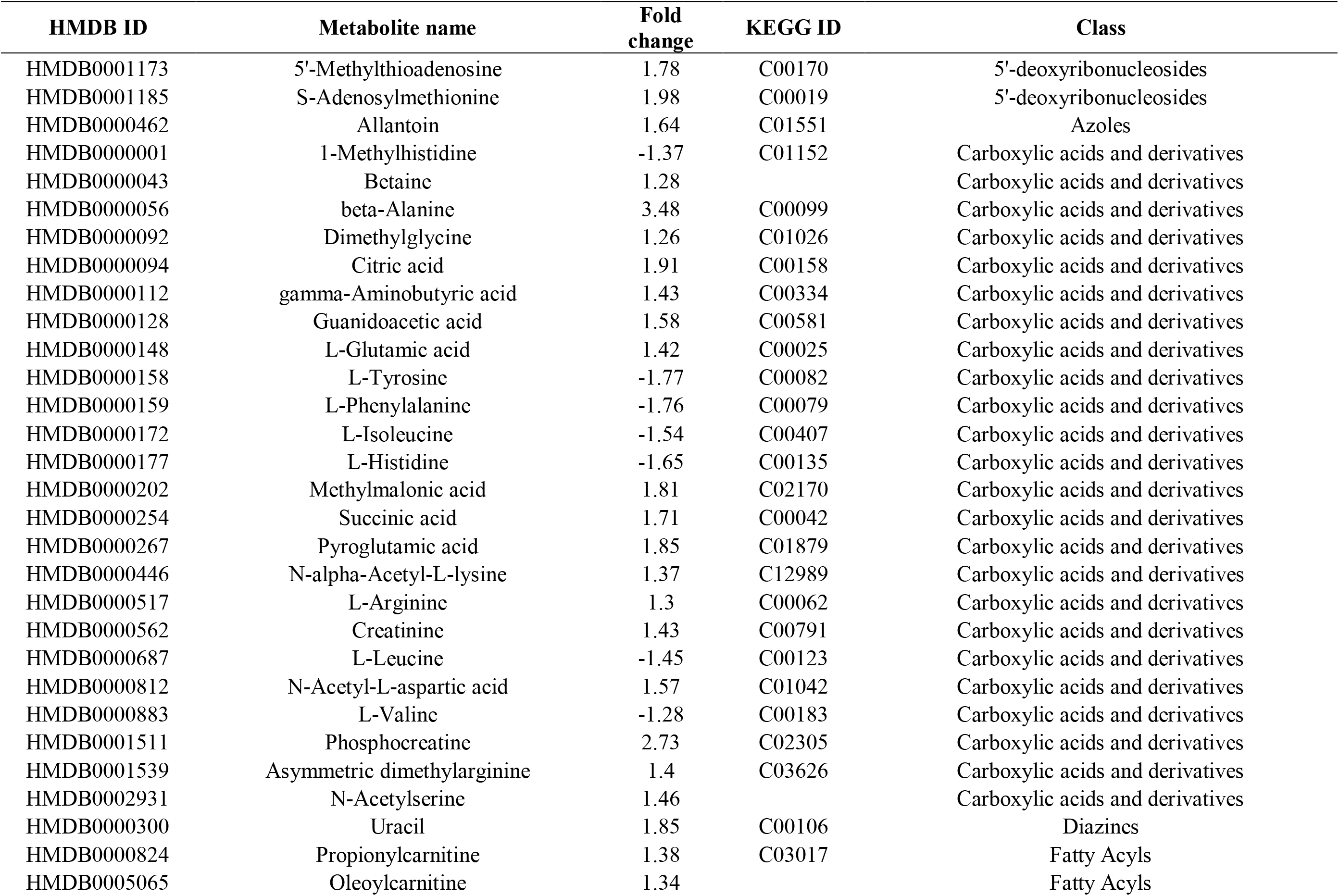
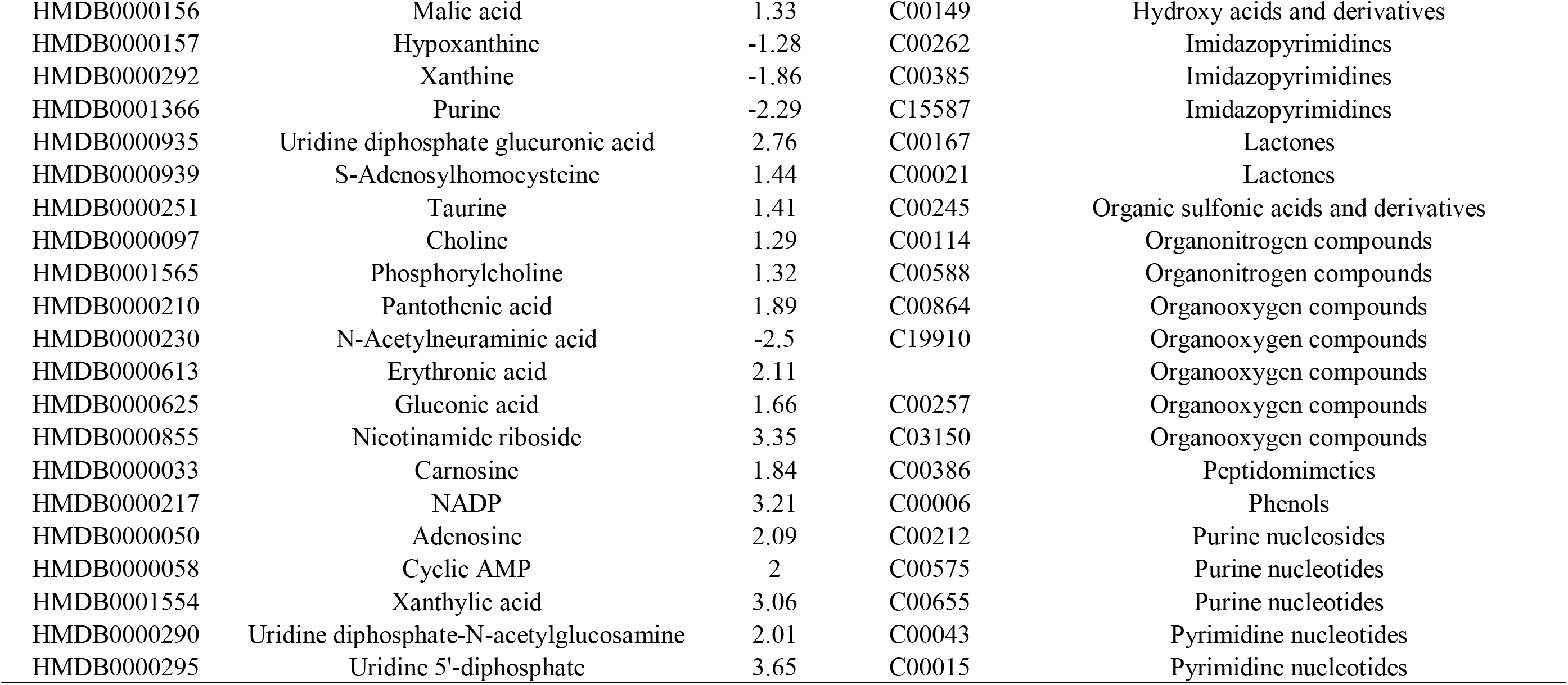
Reference datasets of M8toD6.

| HMDB ID | Metabolite name | Fold change | KEGG ID | Class |
| --- | --- | --- | --- | --- |
| HMDB0001173 | 5'-Methylthioadenosine | 1.78 | C00170 | 5'-deoxyribonucleosides |
| HMDB0001185 | S-Adenosylmethionine | 1.98 | C00019 | 5'-deoxyribonucleosides |
| HMDB0000462 | Allantoin | 1.64 | C01551 | Azoles |
| HMDB0000001 | 1-Methylhistidine | -1.37 | C01152 | Carboxylic acids and derivatives |
| HMDB0000043 | Betaine | 1.28 |  | Carboxylic acids and derivatives |
| HMDB0000056 | beta-Alanine | 3.48 | C00099 | Carboxylic acids and derivatives |
| HMDB0000092 | Dimethylglycine | 1.26 | C01026 | Carboxylic acids and derivatives |
| HMDB0000094 | Citric acid | 1.91 | C00158 | Carboxylic acids and derivatives |
| HMDB0000112 | gamma-Aminobutyric acid | 1.43 | C00334 | Carboxylic acids and derivatives |
| HMDB0000128 | Guanidoacetic acid | 1.58 | C00581 | Carboxylic acids and derivatives |
| HMDB0000148 | L-Glutamic acid | 1.42 | C00025 | Carboxylic acids and derivatives |
| HMDB0000158 | L-Tyrosine | -1.77 | C00082 | Carboxylic acids and derivatives |
| HMDB0000159 | L-Phenylalanine | -1.76 | C00079 | Carboxylic acids and derivatives |
| HMDB0000172 | L-Isoleucine | -1.54 | C00407 | Carboxylic acids and derivatives |
| HMDB0000177 | L-Histidine | -1.65 | C00135 | Carboxylic acids and derivatives |
| HMDB0000202 | Methylmalonic acid | 1.81 | C02170 | Carboxylic acids and derivatives |
| HMDB0000254 | Succinic acid | 1.71 | C00042 | Carboxylic acids and derivatives |
| HMDB0000267 | Pyroglutamic acid | 1.85 | C01879 | Carboxylic acids and derivatives |
| HMDB0000446 | N-alpha-Acetyl-L-lysine | 1.37 | C12989 | Carboxylic acids and derivatives |
| HMDB0000517 | L-Arginine | 1.3 | C00062 | Carboxylic acids and derivatives |
| HMDB0000562 | Creatinine | 1.43 | C00791 | Carboxylic acids and derivatives |
| HMDB0000687 | L-Leucine | -1.45 | C00123 | Carboxylic acids and derivatives |
| HMDB0000812 | N-Acetyl-L-aspartic acid | 1.57 | C01042 | Carboxylic acids and derivatives |
| HMDB0000883 | L-Valine | -1.28 | C00183 | Carboxylic acids and derivatives |
| HMDB0001511 | Phosphocreatine | 2.73 | C02305 | Carboxylic acids and derivatives |
| HMDB0001539 | Asymmetric dimethylarginine | 1.4 | C03626 | Carboxylic acids and derivatives |
| HMDB0002931 | N-Acetylserine | 1.46 |  | Carboxylic acids and derivatives |
| HMDB0000300 | Uracil | 1.85 | C00106 | Diazines |
| HMDB0000824 | Propionylcarnitine | 1.38 | C03017 | Fatty Acyls |
| HMDB0005065 | Oleoylcarnitine | 1.34 |  | Fatty Acyls |
| HMDB0000156 | Malic acid | 1.33 | C00149 | Hydroxy acids and derivatives |
| HMDB0000157 | Hypoxanthine | -1.28 | C00262 | Imidazopyrimidines |
| HMDB0000292 | Xanthine | -1.86 | C00385 | Imidazopyrimidines |
| HMDB0001366 | Purine | -2.29 | C15587 | Imidazopyrimidines |
| HMDB0000935 | Uridine diphosphate glucuronic acid | 2.76 | C00167 | Lactones |
| HMDB0000939 | S-Adenosylhomocysteine | 1.44 | C00021 | Lactones |
| HMDB0000251 | Taurine | 1.41 | C00245 | Organic sulfonic acids and derivatives |
| HMDB0000097 | Choline | 1.29 | C00114 | Organonitrogen compounds |
| HMDB0001565 | Phosphorylcholine | 1.32 | C00588 | Organonitrogen compounds |
| HMDB0000210 | Pantothenic acid | 1.89 | C00864 | Organooxygen compounds |
| HMDB0000230 | N-Acetylneuraminic acid | -2.5 | C19910 | Organooxygen compounds |
| HMDB0000613 | Erythronic acid | 2.11 |  | Organooxygen compounds |
| HMDB0000625 | Gluconic acid | 1.66 | C00257 | Organooxygen compounds |
| HMDB0000855 | Nicotinamide riboside | 3.35 | C03150 | Organooxygen compounds |
| HMDB0000033 | Carnosine | 1.84 | C00386 | Peptidomimetics |
| HMDB0000217 | NADP | 3.21 | C00006 | Phenols |
| HMDB0000050 | Adenosine | 2.09 | C00212 | Purine nucleosides |
| HMDB0000058 | Cyclic AMP | 2 | C00575 | Purine nucleotides |
| HMDB0001554 | Xanthylic acid | 3.06 | C00655 | Purine nucleotides |
| HMDB0000290 | Uridine diphosphate-N-acetylglucosamine | 2.01 | C00043 | Pyrimidine nucleotides |
| HMDB0000295 | Uridine 5'-diphosphate | 3.65 | C00015 | Pyrimidine nucleotides |

**Extended Data Fig. 2.**
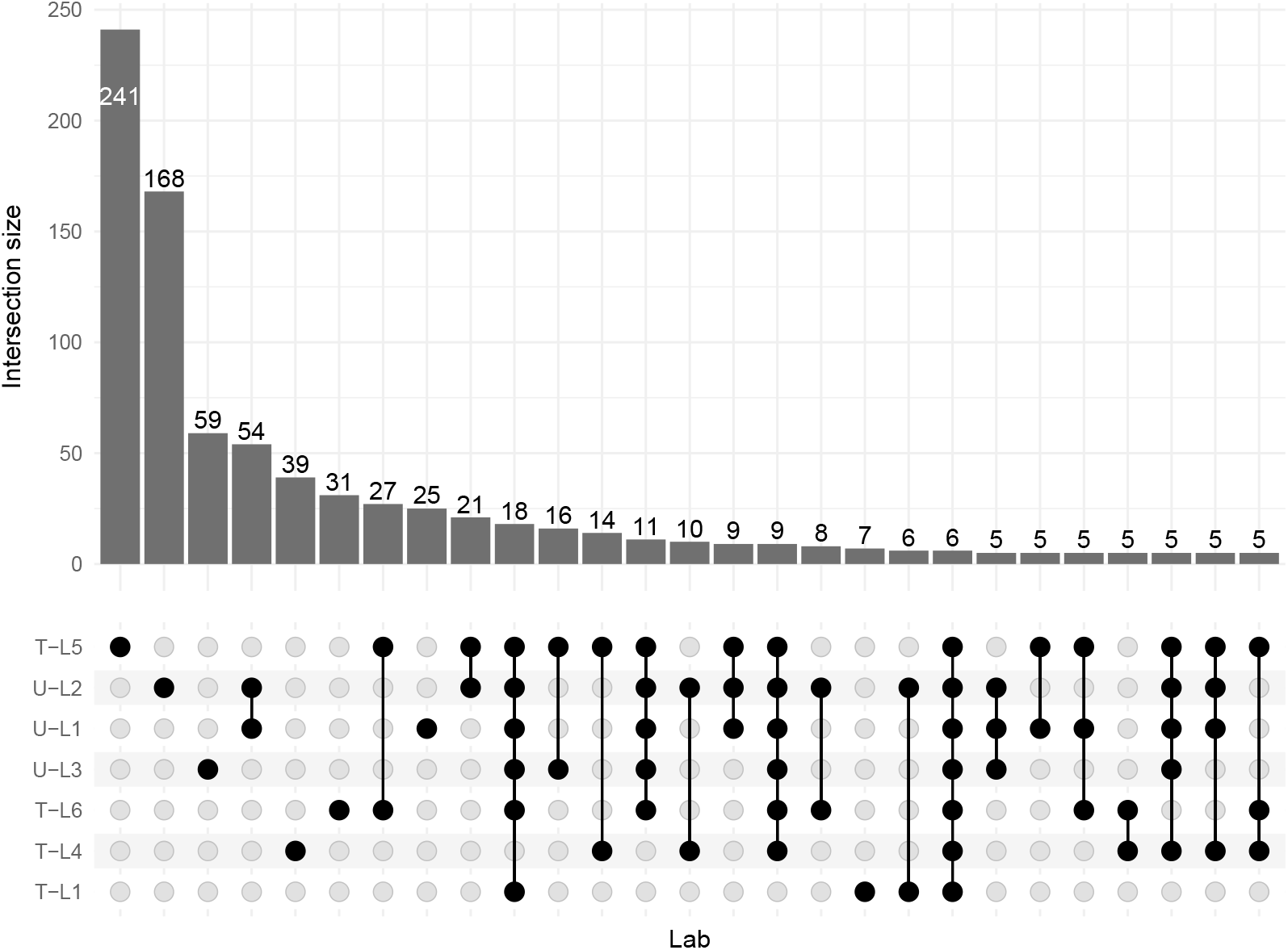
Concordance of detected metabolites among laboratories. The intersection size of detected metabolites among seven datasets generated in different laboratories was shown.

**Extended Data Fig. 3.**
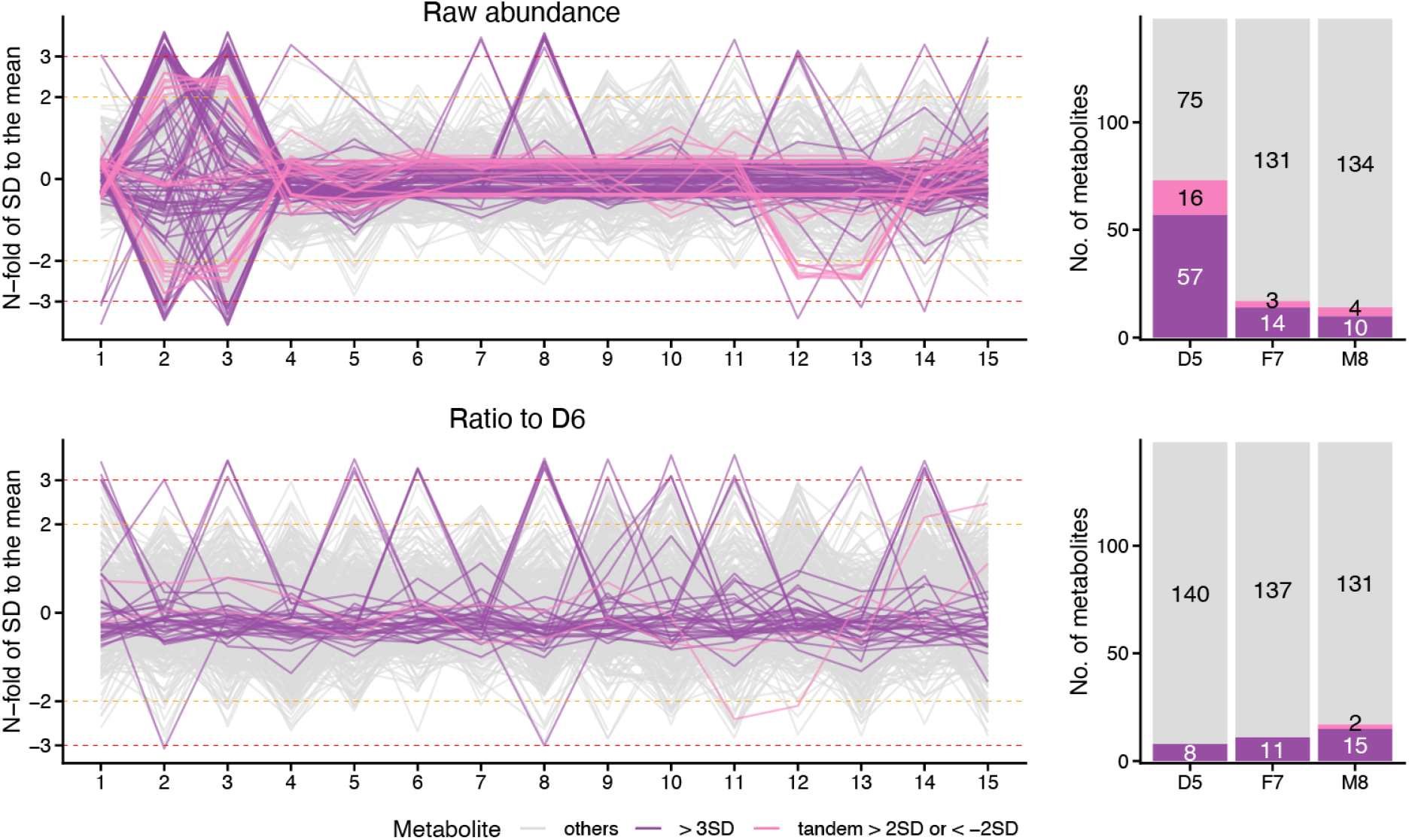
Ratio-based metabolite profiling improves the stability of continuous monitoring of each metabolite measurement. The Levey-Jennings plot of metabolites detected in all 15 batches. Different colors represent different groups of metabolites, indicating systematic deviation > ±3 SD; > ±2 SD, and others.

**Extended Data Fig. 4.**
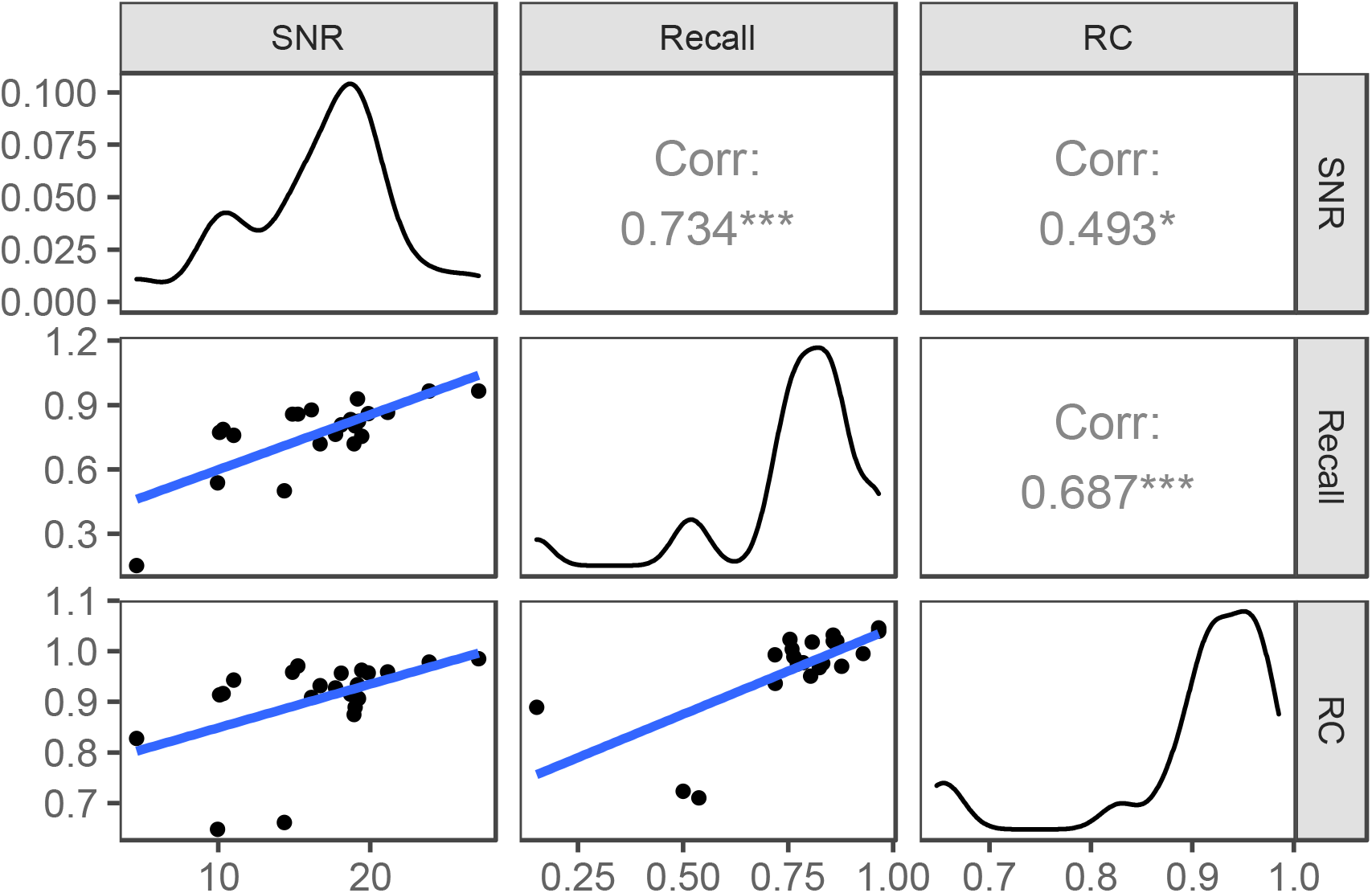
Scatter plot matrices for SNR, Recall and RC.

## References

1. Holmes, E., Wilson, I. D. & Nicholson, J. K. Metabolic Phenotyping in Health and Disease. Cell 134, 714–717 (2008).

2. Nicholson, J. K. et al. Metabolic phenotyping in clinical and surgical environments. Nature 491, 384–392 (2012).

3. Johnson, C. H., Ivanisevic, J. & Siuzdak, G. Metabolomics: beyond biomarkers and towards mechanisms. Nat. Rev. Mol. Cell Biol. 2016 177 17, 451–459 (2016).

4. Wang, T. J. et al. Metabolite profiles and the risk of developing diabetes. Nat. Med. 2011 174 17, 448–453 (2011).

5. Dunn, W. B. et al. Molecular phenotyping of a UK population: defining the human serum metabolome. Metabolomics 11, 9–26 (2015).

6. Liu, J. et al. Metabolomics based markers predict type 2 diabetes in a 14-year follow-up study. Metabolomics 13, 104 (2017).

7. Cirulli, E. T. et al. Profound Perturbation of the Metabolome in Obesity Is Associated with Health Risk. Cell Metab. 29, 488–500 (2019).

8. De Livera, A. M. et al. Statistical Methods for Handling Unwanted Variation in Metabolomics Data. Anal. Chem. 87, 3606–3615 (2015).

9. Lewis, M. R. et al. Development and Application of Ultra-Performance Liquid Chromatography-TOF MS for Precision Large Scale Urinary Metabolic Phenotyping. Anal. Chem. 88, 9004–9013 (2016).

10. Wehrens, R. et al. Improved batch correction in untargeted MS-based metabolomics. Metabolomics 12, 88 (2016).

11. Viant, M. R. et al. Use cases, best practice and reporting standards for metabolomics in regulatory toxicology. Nat. Commun. 10, 3041 (2019).

12. Lippa, K. A. et al. Reference materials for MS-based untargeted metabolomics and lipidomics: a review by the metabolomics quality assurance and quality control consortium (mQACC). Metabolomics 18, 24 (2022).

13. Blaise, B. J. et al. Statistical analysis in metabolic phenotyping. Nat. Protoc. 16, 4299–4326 (2021).

14. Kim, T. et al. A hierarchical approach to removal of unwanted variation for large-scale metabolomics data. Nat. Commun. 12, 4992 (2021).

15. ISO Guide 33:2015. Reference Materials—Good Practice in Using Reference Materials. vol. 2015 (International Organization for Standardization, Geneva, Switzerland., 2015).

16. Telu, K. H., Yan, X., Wallace, W. E., Stein, S. E. & Simõn-Manso, Y. Analysis of human plasma metabolites across different liquid chromatography/mass spectrometry platforms: Cross-platform transferable chemical signatures. Rapid Commun. Mass Spectrom. 30, 581–593 (2016).

17. Beger, R. D. et al. Towards quality assurance and quality control in untargeted metabolomics studies. Metabolomics 15, 4 (2019).

18. Triebl, A. et al. Shared reference materials harmonize lipidomics across MS-based detection platforms and laboratories. J. Lipid Res. 61, 105–115 (2020).

19. Dunn, W. B. et al. Quality assurance and quality control processes: summary of a metabolomics community questionnaire. Metabolomics 13, 50 (2017).

20. Bearden, D. W. et al. The New Data Quality Task Group (DQTG): ensuring high quality data today and in the future. Metabolomics 10, 539–540 (2014).

21. Lindon, J. C. et al. Summary recommendations for standardization and reporting of metabolic analyses. Nat. Biotechnol. 23, 833–838 (2005).

22. Phinney, K. W. et al. Development of a standard reference material for metabolomics research. Anal. Chem. 85, 11732–11738 (2013).

23. Simón-Manso, Y. et al. Metabolite profiling of a NIST standard reference material for human plasma (SRM 1950): GC-MS, LC-MS, NMR, and clinical laboratory analyses, libraries, and web-based resources. Anal. Chem. 85, 11725–11731 (2013).

24. Bearden, D. W. et al. Metabolomics test materials for quality control: A study of a urine materials suite. Metabolites 9, 270 (2019).

25. Aristizabal-Henao, J. J., Jones, C. M., Lippa, K. A. & Bowden, J. A. Nontargeted lipidomics of novel human plasma reference materials: hypertriglyceridemic, diabetic, and African-American. Anal. Bioanal. Chem. 412, 7373–7380 (2020).

26. Aristizabal-Henao, J. J. et al. Metabolomic Profiling of Biological Reference Materials using a Multiplatform High-Resolution Mass Spectrometric Approach. J. Am. Soc. Mass Spectrom. 32, 2481–2489 (2021).

27. Siskos, A. P. et al. Interlaboratory Reproducibility of a Targeted Metabolomics Platform for Analysis of Human Serum and Plasma. Anal. Chem. 89, 656–665 (2017).

28. Townsend, M. K. et al. Reproducibility of Metabolomic Profiles among Men and Women in 2 Large Cohort Studies. Clin. Chem. 59, 1657–1667 (2013).

29. Zhang, X., Dong, J. & Raftery, D. Five Easy Metrics of Data Quality for LC-MS-Based Global Metabolomics. Anal. Chem. 92, 12925–12933 (2020).

30. Koo, T. K. & Li, M. Y. A Guideline of Selecting and Reporting Intraclass Correlation Coefficients for Reliability Research. J. Chiropr. Med. 15, 155–163 (2016).

31. Sampson, J. N. et al. Metabolomics in epidemiology: Sources of variability in metabolite measurements and implications. Cancer Epidemiol. Biomarkers Prev. 22, 631–640 (2013).

32. Evans, A. M. et al. Dissemination and analysis of the quality assurance (QA) and quality control (QC) practices of LC–MS based untargeted metabolomics practitioners. Metabolomics 16, 113 (2020).

33. Want, E. J. et al. Global metabolic profiling procedures for urine using UPLC-MS. Nat. Protoc. 5, 1005–1018 (2010).

34. Dunn, W. B. et al. Procedures for large-scale metabolic profiling of serum and plasma using gas chromatography and liquid chromatography coupled to mass spectrometry. Nat. Protoc. 6, 1060–1083 (2011).

35. Broadhurst, D. et al. Guidelines and considerations for the use of system suitability and quality control samples in mass spectrometry assays applied in untargeted clinical metabolomic studies. Metabolomics 14, 72 (2018).

36. Alseekh, S. et al. Mass spectrometry-based metabolomics: a guide for annotation, quantification and best reporting practices. Nat. Methods 18, 747–756 (2021).

37. Sánchez-Illana, Á. et al. Evaluation of batch effect elimination using quality control replicates in LC-MS metabolite profiling. Anal. Chim. Acta 1019, 38–48 (2018).

38. Bowden, J. A. et al. Harmonizing lipidomics: NIST interlaboratory comparison exercise for lipidomics using SRM 1950–Metabolites in Frozen Human Plasma. J. Lipid Res. 58, 2275–2288 (2017).

39. Dudzik, D., Barbas-Bernardos, C., Garcaí, A. & Barbas, C. Quality assurance procedures for mass spectrometry untargeted metabolomics. a review. J. Pharm. Biomed. Anal. 147, 149–173 (2018).

40. Izumi, Y. et al. Inter-Laboratory Comparison of Metabolite Measurements for Metabolomics Data Integration. Metabolites 9, 257 (2019).

41. Zheng, Y. et al. Ratio-based multiomic profiling using universal reference materials empowers data integration. Preprint at doi:10.1101/2022.10.24.513612v1.

42. Ren, L. et al. Quartet DNA reference materials and datasets for comprehensively evaluating germline variants calling performance. Preprint at https://doi.org/10.1101/2022.09.28.50984443.

43. Yu, Y. et al. Quartet RNA reference materials and ratio-based reference datasets for reliable transcriptomic profiling. Preprint at https://doi.org/10.1101/2022.09.26.507265

44. Tian, S. et al. Quartet protein reference materials and datasets for multi-platform assessment of label-free proteomics. Preprint at https://doi.org/10.1101/2022.10.25.513670

45. Yu, Y. et al. Correcting batch effects in large-scale multiomic studies using a reference-material-based ratio method. Preprint at https://doi.org/10.1101/2022.10.19.507549

46. Yang, J. et al. The Quartet Data Portal: integration of community-wide resources for multiomics quality control. Preprint at https://doi.org/10.1101/2022.09.26.507202

47. Naz, S., Vallejo, M., García, A. & Barbas, C. Method validation strategies involved in non-targeted metabolomics. J. Chromatogr. A 1353, 99–105 (2014).

48. Wheeler, H. E. & Dolan, M. E. Lymphoblastoid cell lines in pharmacogenomic discovery and clinical translation. Pharmacogenomics 13, 55–70 (2012).

49. Ozgyin, L., Horvath, A., Hevessy, Z. & Balint, B. L. Extensive epigenetic and transcriptomic variability between genetically identical human B-lymphoblastoid cells with implications in pharmacogenomics research. Sci. Rep. 9, 4889 (2019).

50. Cai, Y., Weng, K., Guo, Y., Peng, J. & Zhu, Z. J. An integrated targeted metabolomic platform for high-throughput metabolite profiling and automated data processing. Metabolomics 11, 1575–1586 (2015).

51. Evans, A. M. et al. High Resolution Mass Spectrometry Improves Data Quantity and Quality as Compared to Unit Mass Resolution Mass Spectrometry in High-Throughput Profiling Metabolomics. J. Postgenomics Drug Biomark. Dev. 04, 2 (2014).

